# Genome-wide profiling of DNA repair proteins identifies higher-order coordination in single cells

**DOI:** 10.1101/2023.05.10.540169

**Authors:** Kim L. de Luca, Pim M. J. Rullens, Magdalena A. Karpinska, Sandra S. de Vries, Agnieszka Gacek-Matthews, Lőrinc S. Pongor, Gaëlle Legube, Joanna W. Jachowicz, A. Marieke Oudelaar, Jop Kind

**Affiliations:** Hubrecht Institute, Royal Netherlands Academy of Arts and Sciences (KNAW) & University Medical Center Utrecht; Utrecht, the Netherlands; Oncode Institute; the Netherlands; Department of Molecular Biology, Faculty of Science, Radboud Institute for Molecular Life Sciences, Radboud University; Nijmegen, the Netherlands; Genome Organization and Regulation, Max Planck Institute for Multidisciplinary Sciences; Göttingen, Germany; Institute of Molecular Biotechnology of the Austrian Academy of Sciences (IMBA), Vienna Biocenter (VBC); Vienna, Austria; MCD, Centre de Biologie Intégrative (CBI), CNRS, Université de Toulouse; Toulouse, France; Cancer Genomics and Epigenetics Core Group, Hungarian Center of Excellence for Molecular Medicine (HCEMM); Szeged, Hungary

## Abstract

Accurate repair of DNA damage is critical for maintenance of genomic integrity and cellular viability. Because damage occurs non-uniformly across the genome, single-cell resolution is required for proper interrogation, but sensitive detection has remained challenging. Here, we present a comprehensive analysis of repair protein localization in single cells using DamID and ChIC sequencing techniques. This study reports genome-wide binding profiles in response to DNA double-strand breaks induced by AsiSI, and explores variability in genomic damage locations and associated repair features in the context of spatial genome organization. By unbiasedly detecting repair factor localization, we find that repair proteins often occupy entire topologically associating domains, mimicking variability in chromatin loop anchoring. Moreover, we demonstrate the formation of multi-way chromatin hubs in response to DNA damage. Notably, larger hubs show increased coordination of repair protein binding, suggesting a preference for cooperative repair mechanisms. Together, our work offers new insights into the heterogeneous processes underlying genome stability in single cells.

## Main

The eukaryotic nucleus is constantly exposed to endogenous and exogenous sources of damage to the genome. Among these, double-stranded breaks (DSBs) in the DNA are particularly hazardous lesions because they completely sever the DNA fiber, leaving the genome at risk of small nucleotide changes and larger structural aberrations such as translocations and deletions. These types of genomic instability are associated with tumorigenesis as well as aging-related diseases. To ensure genome integrity, the cell is dependent on the DNA damage response (DDR), an intricate signaling cascade that includes recognition, processing, and restoration of the lesion^1^. The two main groups of DSB repair pathways are end joining (EJ) and homology-directed repair (HDR). EJ involves re-ligation of minimally processed DNA ends and can occur throughout interphase, while HDR requires a homologous template sequence (usually its sister chromatid) and is therefore generally restricted to S and G2 phase^2^. Processing and repair outcome are thus influenced by the phase of the cell cycle, complexity of the DSB, as well as its transcriptional status, chromatin environment, and location within the nucleus^3,4^. In response to damage, DDR proteins accumulate into DNA repair foci, which are formed by local rearrangement of the chromatin at the level of topologically associating domains (TADs)^5^. Further, damaged chromatin exhibits more large-scale mobility, forming clusters of DSBs that reside in specific sub-compartments^6,7^.

Given the variability in the occurrence and repair of DNA damage across cells, it becomes imperative to collect information from individual cells to accurately profile their distribution. Single-cell detection of spontaneous damage was described using whole-genome amplification^8^, but insight into DNA repair at this level has not been reported from a genomic perspective. In this study, we address this technological gap by presenting a detailed analysis of DSB repair factor localization in single cells. We use two methods to map DNA-binding proteins, namely DamID^9^ and ChIC^10^, combined with a new computational framework for signal detection, and compare our approach to the state of the art. Additionally, we report simultaneous measurement of repair protein signal (with DamID) and chromatin features such as histone modifications or structural proteins (with ChIC) in the same cell^11^, enabling direct analysis of the interplay between repair proteins and the damaged chromatin substrate.

A distinctive property of single-cell data lies in the ability to measure signal across the entire genome on a per-cell basis. With a sufficient number of cells, and diversity in signal among those cells, patterns can start to emerge that reflect underlying processes of interest. To illustrate this concept, we induce damage at many (∼100) known locations in the human genome with the DIvA system^12^, and quantify repair protein binding at all sites within individual cells. Specifically, we investigate whether sites are simultaneously occupied by the repair machinery, referred to as “coordination”. We explore such coordination in the context of damage-specific genome reorganization.

Overall, our data reveal heterogeneity in repair factor localization that was previously unappreciated. We demonstrate the utility of (multifactorial) protein profiling in single cells, setting the stage for future investigations into DNA repair and genome stability.

### Detecting double-strand break repair proteins genome-wide with DamID and ChIC

We established an experimental and computational workflow to unbiasedly identify DSB repair profiles in single cells (Fig. 1a). Using DamID^9,13–15^ and ChIC^10,16,17^, we measured genomic contacts of proteins involved in EJ and HDR, choosing different Dam-fusion proteins (for DamID) or antibodies (for ChIC). After filtering on quality criteria (see Methods), we obtained a collective total of ∼15,000 single-cell profiles of DSB repair proteins 53BP1, MDC1, and γH2AX, and HDR-specific proteins RAD51 and BRCA1 (Fig. 1b, Extended Data Fig. 1a-b). We used the previously established human DIvA cell line, which generates DSBs at sequence-specific positions in the genome by the endonuclease AsiSI-ER under control of 4-hydroxytamoxifen (4OHT)^12,18^.

**Fig. 1.**
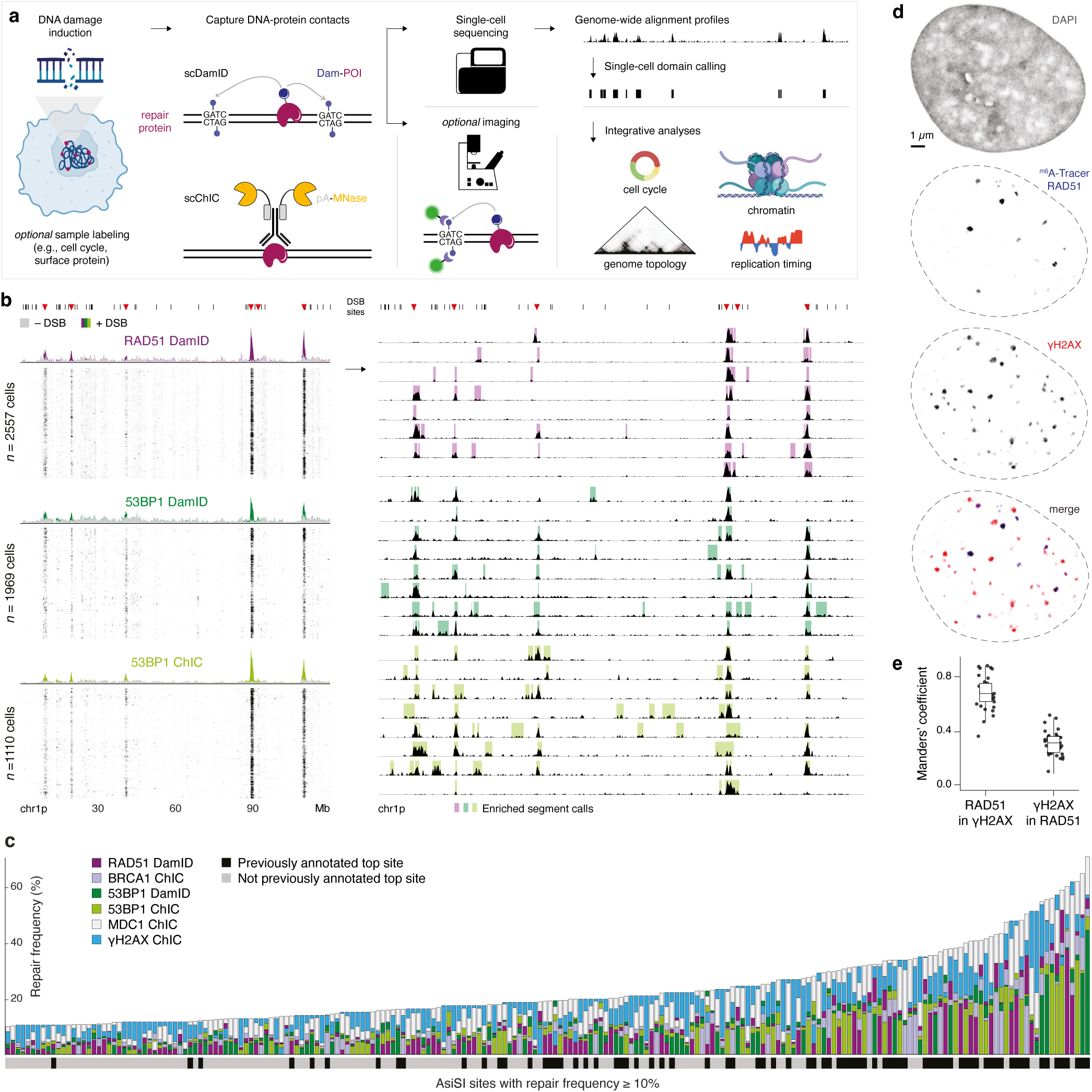
Detecting double-strand break repair proteins with DamID and ChIC. **a,** Overview of experimental setup and integrations. **b**, Signal on chromosome 1p. Left: single-cell heatmaps (RPKM) in +DSB condition, with single-cell aggregates of +DSB (colored) and –DSB (gray) conditions on top. Right: single-cell line plots with overlaid MSR calls. AsiSI motifs, annotated with black lines, and red triangles indicating frequently cleaved (or “top”) sites. **c**, Bar plot of all AsiSI sites with repair protein frequency ≥ 10%, colored by repair protein frequencies per dataset (target protein & method) per site. Sites are ordered (on the x-axis) by increasing absolute repair protein frequency (*i.e.*, highest frequency in any dataset). Per site, bars are ordered by increasing repair protein frequency per dataset (front to back; *i.e*., bars are not stacked). Bottom horizontal bar indicates previous (lack of) annotation as top site. **d**, Confocal images of one representative nucleus showing DAPI, RAD51 DamID m6A-Tracer, and endogenous γH2AX immunofluorescent staining. **e**, Quantification of signal colocalization (Manders’ A and B per nucleus), *n* = 33 nuclei.

To determine significant enrichments of repair occupancy on the genome, we developed a computational method for single-cell domain calling. We modified a multi-scale representation of genomic signals (MSR) approach^19^ that unbiasedly identifies enrichments of variable sizes (Extended Data Fig. 2, Methods). Upon damage induction, specific repair signal accumulates at DSB sites (Fig. 1b, Extended Data Fig. 1c-e), which compares well with publicly available population-based ChIP-seq datasets (Extended Data Fig. 1e-g). Measured across all cells and DSB sites, we calculated the proportion of cells in which each DSB was bound by the different target proteins (referred to as “repair protein frequency”). Our collective dataset indicates superior sensitivity compared to previous reports (Fig. 1c), allowing for the study of a broader range of infrequently captured DSB sites. In accordance with the DDR signaling cascade, nearly all DSB sites are most frequently occupied by mediator of DNA damage checkpoint protein 1 (MDC1) and ubiquitous DSB marker γH2AX (the direct chromatin substrate for MDC1 binding), followed by EJ-promoting 53BP1, and HDR-specific RAD51 and BRCA1 (Fig. 1c). Nonetheless, repair signatures clearly vary across sites, distinguishing between EJ and HDR preference. To further quantify the repair domains per cell, we defined a set of single-cell profiles in which we could most accurately determine the number of MSR segments, also referred to as repair protein domains (see Methods). At the median sequencing depth (3×10^3^ UMIs), DSB-treated and control cells respectively contained 28 and 3 repair-enriched segments, reaching a plateau at 50 and 7 segments (Extended Data Fig. 1h), in agreement with the number of AsiSI-induced 53BP1 foci detected by imaging^20^.

DSB repair has been widely studied with imaging-based methods that detect proteins, while DamID relies on detection of genomic DNA that has been marked by the Dam-fusion protein. We previously developed the m6A-Tracer system for visualization and tracking of Dam-methylated contacts^21^, and apply it here to validate that HDR-specific Dam-RAD51 generates foci that visually overlap with γH2AX, a universal DSB marker (Fig. 1d). Of the DamID RAD51 signal, ∼65% colocalizes with γH2AX; vice versa, a smaller ∼30% of γH2AX signal corresponds to Dam-RAD51 (Fig. 1e). The latter is anticipated to be lower since not all γH2AX-marked DSBs are repaired by RAD51, as can also be observed from the smaller number of Dam-RAD51 foci.

Together, these results show that implementation of DamID and ChIC can be used for sensitive and specific single-cell genomics as well as quantitative imaging analyses of DSB repair.

### Homology-directed repair mediated by RAD51 correlates with replication timing and transcriptional activity

Repair pathway choice between EJ and HDR is highly regulated at multiple levels, including nuclear structure, global spatial genome organization, local chromatin context, and sequence specificity^22^. In addition to such regulatory processes, the cell cycle state of individual cells has been linked to heterogeneity in repair pathway usage^2,23^. Our DamID experimental setup includes recording of live-cell DNA content during fluorescence-activated cell sorting (FACS), thereby establishing a procedure to address DSB repair protein occupancy in relation to the cell cycle at unprecedented resolution (Fig. 2a). We observed that some DSB sites exhibit differences in repair protein frequency according to cell cycle phase, particularly when bound by HDR factors (Extended Data Fig. 3a). To further explore this relationship, we ordered all Dam-RAD51 cells on their cell cycle stage, and noticed differences in repair enrichment at DSB sites during S phase (Fig. 2b-c), a genome-wide trend that is not present for 53BP1 (Extended Data Fig. 3b). Because HDR requires a sister chromatid as its repair template, we annotated the genomic regions by their replication timing (RT) using publicly available Repli-Seq data^24,25^. This showed considerable concordance between RT of a site and the relative frequency with which it is bound by Dam-RAD51 along S and G2 (Fig. 2d), in agreement with imaging data suggesting that active replication influences HDR employment^23^.

**Fig. 2.**
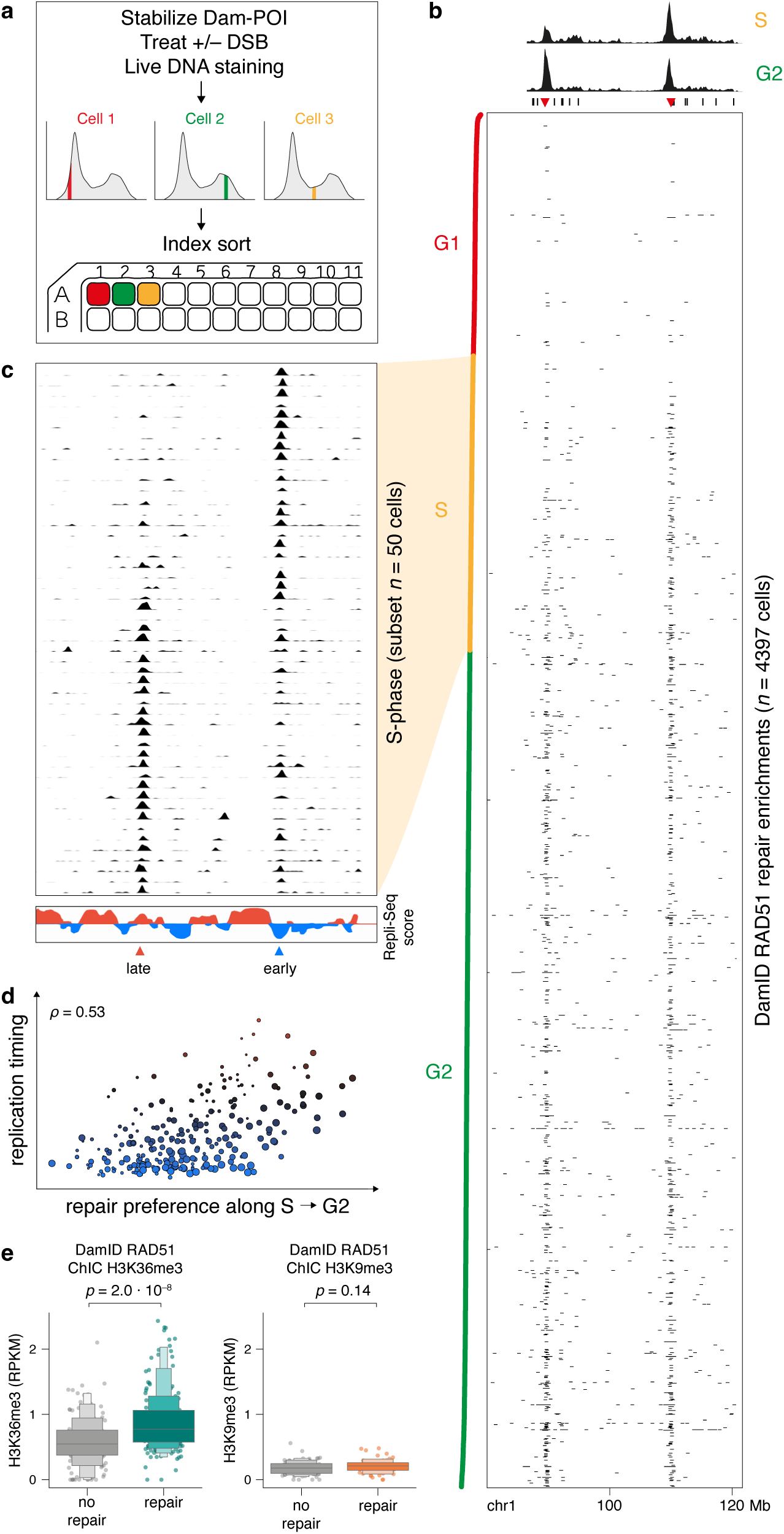
RAD51 correlates with cell cycle phase, replication timing, and transcriptionally active chromatin in single cells. **a**, Overview of DamID experimental setup up to and including FACS. **b**, Heatmap showing binarized RAD51 DamID enrichment (MSR calls) of 4397 single cells, at a selected region on chromosome 1. DNA content as measured by live Hoechst staining during FACS is annotated along the vertical axis (left side): G1 in red, S in yellow, G2 in green. Single-cell aggregates of +DSB condition for S and G2 phases on top, with AsiSI motifs annotated as in Fig. 1b. **c**, Line plots showing RAD51 signal (RPKM) for a subset of cells in S phase. **d**, Quantification of correlation between cell cycle preference and Repli-Seq scores for AsiSI clusters with repair protein frequency ≥ 5%. Size of circles indicates repair protein frequency. Color of circles corresponds to the *y*-axis. **e**, Boxen plots showing H3K36me3 (left) and H3K9me3 (right) signal (RPKM) at AsiSI site 505, for cells with (right) and without (left) RAD51 enrichment. Statistical significance was calculated by two-sample Kolmogorov-Smirnov (KS) test.

RT is significantly correlated with chromatin state and genome organization^26,27^; moreover, RT-driven remodeling of the epigenome has been linked to cancer-specific chromosomal rearrangements^28^. Several studies have further highlighted the role of chromatin in DNA repair^29–38^; notably, RAD51 is preferentially recruited to transcribed loci enriched in active histone modification H3K36me3^29,39^. We sought to directly measure the relationship between DNA repair and chromatin, by jointly profiling RAD51 occupancy (with DamID) and either H3K36me3 or repressive mark H3K9me3 (with ChIC, using an antibody) in the same cell (see Methods and ^11^).

Combined single-cell Dam&ChIC profiles were of similar quality to DamID as the only genomic readout (Extended Data Fig. 3c-d), and ChIC histone modification signal was in excellent concordance with published ChIP-seq (Extended Data Fig. 3e-f). We show that H3K36me3 enrichment is significantly higher in cells that have RAD51 binding than in cells without, for a given AsiSI site (Fig. 2e) and across all sites (Extended Data Fig. 3g). In contrast, H3K9me3 is expectedly low, and similar between bound states (Fig. 2e, Extended Data Fig. 3g). This corroborates, at the single-cell level, the previously established link between HDR and transcriptionally active chromatin.

### Repair protein occupancy in single cells follows pre-existing genome topology

Next, we further explored the binding of repair proteins at DSBs, and the potential “spreading” of proteins along the genome. In particular, we set out to address chromatin conformation of DSB repair sites at the single-cell level. Recent population-based work has reported local chromatin reorganization upon DNA damage^5,7^, and proposed cohesin-mediated loop extrusion as the mechanism by which DNA repair domains are established^5^. This further substantiates the thought that chromatin is compacted during the DDR, forming incontiguous domains according to topological features^40–44^.

#### Repair factors are constrained within topologically associating domains in multiple modalities

To examine the interplay between genome topology and repair factor binding in single cells, we projected our DamID 53BP1-enriched segments onto Hi-C maps (Extended Data Fig. 4a, Fig. 3a). Repair domain enrichment occurs in different scenarios, varying mostly according to extent, directionality, and “modality” of spreading. Modality refers to whether one (*i.e.*, unimodal) or different (*i.e.*, multimodal) enrichment patterns are found, and is thus a measure of intercellular variability. Domains of repair occupancy seem generally constrained by borders of chromosomal domains. In highly insulated chromatin, 53BP1 segments clearly overlap with the local genome structure, while DSB sites devoid of detectable structure show repair occupancy in a scattered fashion, without topological constraints. Of note, cell-to-cell heterogeneity is strongly reflected in the repair protein segments, since a spectrum of domain sizes can be observed (Extended Data Fig. 4a, Fig. 3a). We note that these heterogeneous measurements are a direct result of detecting enrichment without any a priori information of segment size, enabled by our newly implemented multi-scale domain calling approach.

**Fig. 3.**
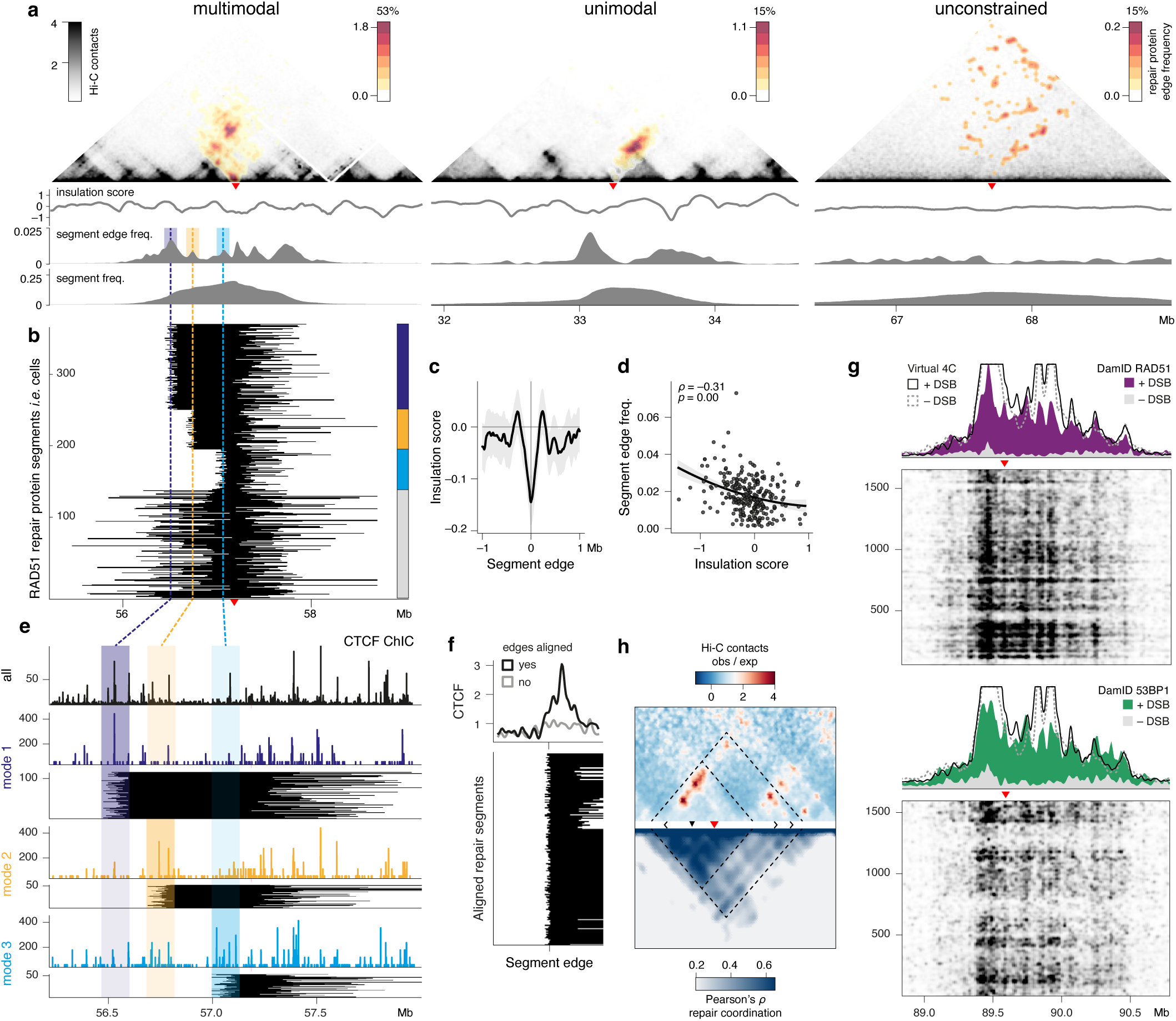
Repair protein spreading in single cells mimics underlying genome topology. **a**, Overlay of 53BP1 DamID repair protein segment edges onto the Hi-C contact matrix for regions harboring a DSB site (red triangle). Repair protein binding signal is colored by the frequencies with which those edges are observed across single cells. The total frequency of each locus (in % of cells bound by repair protein) is indicated above. One example region is shown for different scenarios of repair protein spreading: multimodal (left), unimodal (middle), and unconstrained (right). Dashed colored lines indicate local maxima of edge frequency to the left of the top site: outermost (purple), middle (orange) and nearest (blue) segment edges. Shaded area indicates the window (150 kb) used to group segment edges into “modes”. **b**, Individual repair protein segments as called per single cell, grouped into modes on the left border, based on windows as in **a**. Colored blocks on the right indicate the modes, and all other segments not assigned to any mode. **c**, Quantification of insulation score across all segment edges (in a window of 2 Mb). Shaded area indicates 95% CI. **d**, Quantification (Pearson’s *r*) of repair-based segment edge frequency and topology-based insulation score at that edge. **e**, Co-measurement of Dam-RAD51 and CTCF in the same cells. Upper track is the RPKM-normalized CTCF signal of all cells. Below are three groups of spreading modes as defined in **b**. For each mode, repair protein segments are plotted in the binary lines, and the corresponding CTCF signal of those same cells is plotted in the colored tracks above. Shaded areas indicate spreading modes. **f**, (bottom) Genome-wide version of the locus-specific plots in **e**. All DSB segment edges are aligned and centered, *i.e.*, all colored blocks. (top) The average CTCF signal of those same “aligned” cells (black line corresponds to all colored blocks) and for cells in which no segment edge is found at those same sites (grey line corresponds to all grey blocks). **g**, Comparison of virtual 4C (representation of Hi-C contacts formed along the linear genome) and Dam-RAD51(top) and Dam-53BP1 (bottom) signal. Single-cell aggregates for both conditions are plotted above single-cell heatmaps +/–DSB. h, Comparison of the normalized Hi-C contact matrix (top) and single-cell repair coordination (bottom) for a 5-Mb region. Repair coordination across cells is measured per pair of genomic bins.

To further illustrate multimodal enrichment, we grouped the repair protein segments according to their three most frequent edge positions (Fig. 3a-b). We noticed that 53BP1 segment edges correspond to relative dips in the insulation score, which is a measure of TAD border strength. To investigate if the variability in repair protein spreading is linked to topological variability, we generated split-pool recognition of interactions by tag extension (SPRITE) data. SPRITE is a method that, unlike Hi-C, detects higher-order chromatin structures^45^, which can be interpreted as single-molecule topologies. Indeed, we detect various SPRITE clusters that overlap the three most frequent repair protein segment edges (Extended Data Fig. 4b), mimicking the multiple modes of repair spreading.

Next, we sought to generalize this apparent relationship between repair protein spreading in single cells and topological domains. Accordingly, taken across all DSB sites genome-wide, there is a strong minimum in the insulation score at repair protein segment boundaries, indicating overlap with TAD borders (Fig. 3c). Repair protein segments also more frequently end at stronger borders (*i.e.*, with lower insulation scores), as quantified by the negative overall correlation (Fig. 3d).

#### Structural protein CTCF demarcates individual repair domains

TAD-like partitioning of repair domains is thought to stabilize chromatin topology and thereby safeguard genome integrity^41^. Further, insulator protein CTCF was found to be in close spatial proximity to radiation-induced γH2AX repair foci (as measured by super-resolution microscopy^40^). CTCF-bound loop anchors are also particularly fragile sites (as measured by mapping DSBs^46^). Yet, a genome-centered view of repair factors and 3D organization is lacking. We used the combined Dam&ChIC method to gather genome-wide information of repair protein RAD51 (with DamID) and structural protein CTCF (with ChIC) within the same cell, akin to two-color imaging.

As quality control, alignment of the aggregated CTCF signal on CTCF binding motifs indicates high specificity (Extended Data Fig. 4b), while retaining sensitivity of median ∼10^3^ reads per cell (Extended Data Fig. 4c).

In the combined RAD51&CTCF dataset, we again grouped the repair protein segments (that is, cells) into “modes” based on the RAD51 edge positions. This was done by calling peaks on the distribution of segment edge frequencies (as seen in Fig. 3a) and selecting segments within a 100-kb window surrounding each peak (shaded areas). For each spreading mode, we then plotted the CTCF signal corresponding to the cells in that mode (Fig. 3e, colored lines). By visual inspection, binding of CTCF is consistently highest in the area surrounding its corresponding segment edges. Seemingly high-occupancy CTCF loci (e.g., blue shaded area) may be near-absent in most cells (modes 1 and 2), while particularly enriched in others (mode 3).

We quantify this observation genome-wide, by aligning all repair protein segment edges found within peak mode windows (*i.e.*, all colored blocks across the whole genome), and comparing them with all other segments not found within those windows, (*i.e.*, all grey blocks). CTCF signal is strongly enriched on the aligned segment edges, while no such pattern can be found for unaligned segments (Fig. 3f). With this, our data show that structural proteins such as CTCF demarcate repair domains, further assigning functionality to TAD-like chromatin structures. The observation of heterogeneity within the distributions of repair domains—and their relationship to genome organization—thereby highlights the importance of sequencing-based profiling at single-cell resolution.

### Repair protein signal is highly coordinated within loop extrusion borders

In the comparison with Hi-C above, we defined repair protein occupancy as the enrichment from start to end of the MSR segments. To further strengthen the notion that such occupancy occurs in 3D, we compared repair spreading at a given DSB site to the chromosomal conformation of that locus (“virtual 4C” from Hi-C data). The quantitative repair counts—*i.e.*, sequencing read output, not domain calls—captured by both DamID and ChIC strongly resemble topological contacts along the linear genome (Fig. 3g). Hence, the quantitative in silico population signal, single-cell profiles, and binarized MSR segments all show that DNA repair occurs in the context of genome organization.

Importantly, single-cell Hi-C and super-resolution chromatin tracing methods have indicated that boundaries of TAD-like domains vary across cells, despite preferential anchoring at population-based boundaries^47–50^. We reasoned that, if repair proteins spread according to these topology-driven rules of variability and boundary anchoring, loci with stronger Hi-C contacts should more frequently show coordinated occupancy of repair proteins across single cells. Conversely, repair signal should be independent for loci with weak (or few) Hi-C contacts. Coordination is thus a measure of how frequently a repair protein occupies a DSB locus, measured across all cells: it should reflect the diversity of topological configurations commonly observable in a given Hi-C window. To best interpret chromatin contacts, we calculated the normalized Hi-C matrix (observed/expected), where distinct architectural stripes and dots can be observed (Fig. 3h), which are features of cohesin-mediated loop extrusion. Indeed, we find that repair protein coordination surrounding a DSB corresponds to those features, suggesting that the repair machinery spreads within various preferentially anchored but dynamic single-cell loops (Fig. 3h).

Altogether, we interpret these results to mean that the spreading of repair proteins on the genome follows underlying topology, explaining differences in repaired genome segment size as well as quantitative repair signal across DSB loci.

### Multi-way coordination of repair protein binding at long-range contacts

Besides local reorganization of the genome in response to damage as described in Figure 3, DSBs exhibit intra-nuclear motion on a more global scale^6,20,51–53^. The phenomenon of repair foci clustering was first observed by microscopy^54^: over time, a reduction in the number of γH2AX foci and an increase in their size supported the “breakage-first” theory of chromosomal translocations. Live-cell imaging further showed fusion of separate repair foci by fluorescent tagging of repair proteins 53BP1^20,55,56^ and Rad52^53^. On the genomic (rather than the protein) level, high-throughput sequencing experiments showed that DSB loci physically interact, by comparing chromosomal contacts in damaged and undamaged conditions^6,7,56^. However, no experimental evidence has been presented that directly couples repair foci clustering and genomic identity of DNA breaks.

#### Coordination of single-cell repair protein binding corresponds to Hi-C contact frequency

First, we re-analyzed recent Hi-C data^5^ for downstream visualization and statistical purposes, confirming that some DSB sites form long-range and often inter-chromosomal contacts upon damage (Fig. 4a, Extended Data Fig. 5a). Damage-specific contacts were identified by computing a differential Hi-C matrix, which quantifies the fold-change between control and damage-induced conditions (Fig. 4bi, different *x*-axis region in Fig. 4bii). We defined pairs of sites as “contacting” by setting a threshold on the differential matrix (Extended Data Fig. 5b), and validate Hi-C signal up to +/- 0.5 Mb surrounding those DSB sites, while non-contacting sites are fully devoid (Fig. 4c top).

**Fig. 4.**
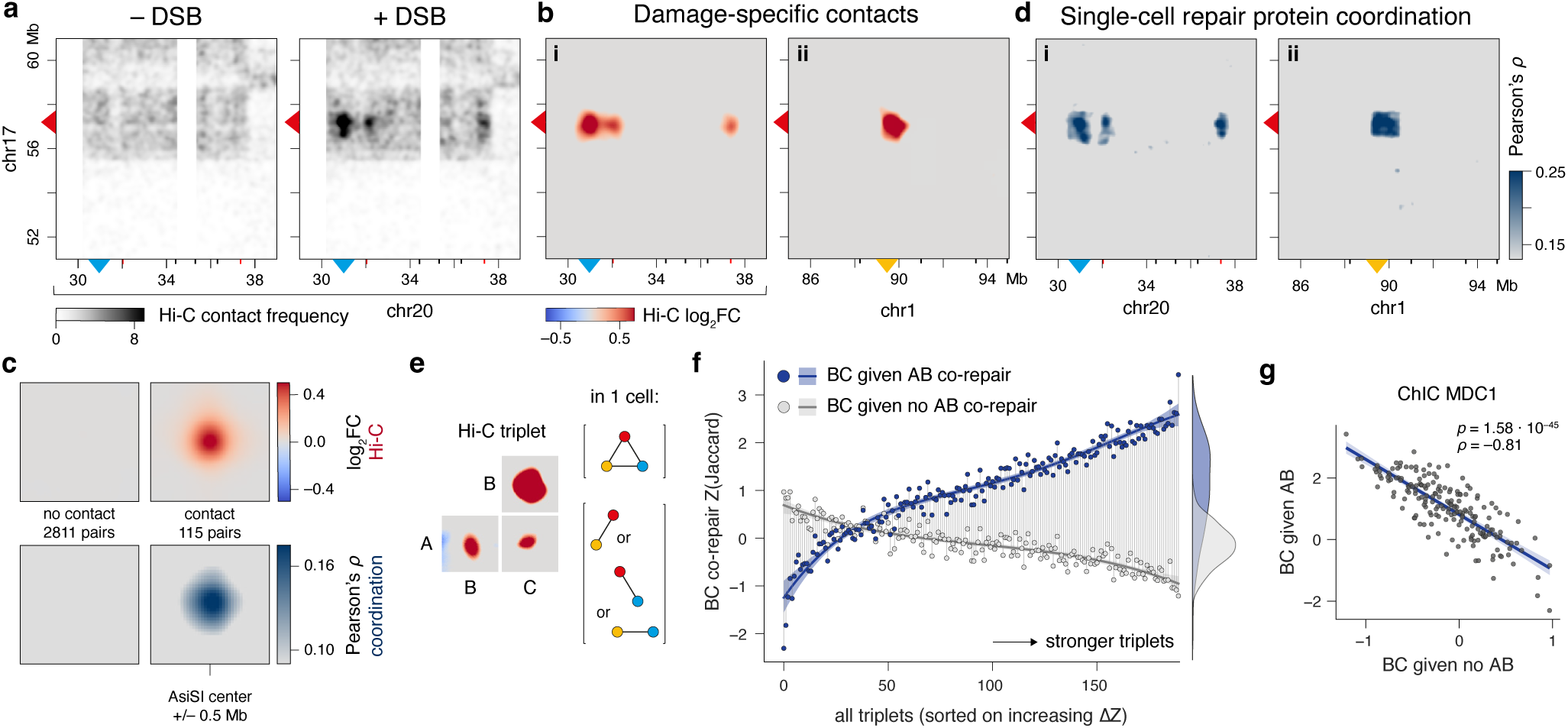
Pairwise repair coordination of long-range contacts in single cells. **a**, Hi-C signal in 10-Mb regions harboring DSB sites. The left and right panel show Hi-C contacts of chr17 (*y*) and chr20 (*x*) in –DSB and +DSB conditions, respectively. **b**, Left panel shows differential Hi-C contacts (log2FC (+DSB / –DSB)) of chr17 with chr20 as presented in **a**. Right panel shows differential Hi-C contacts of a different region between chr17 and chr1. **c**, Comparison of Hi-C contact score (top) and single-cell repair coordination (bottom) across all pairs of DSB sites, categorized by contact (right) or no contact (left) based on log_2_FC threshold. A region of 1 Mb surrounding the AsiSI motif is shown. Coordination is measured as Pearson’s correlation between 40-kb bins across cells. **d**, Single-cell repair coordination (53BP1) of all pairwise combinations of bins in selected regions as in b. Coordination is measured as Pearson’s correlation between 40-kb bins across cells. **e**, Left panel shows Hi-C based definition of a triplet where all three sites A, B and C have pairwise contacts. Right panel shows a schematic representation of higher-order (top) and mutually exclusive (bottom) contacts. **f**, Scatter plot measures three-way coordination of triplets. Each triplet is represented twice: BC co-repair is quantified as the *Z*-score normalized Jaccard index between B and C in 1) cells that have AB co-repair (blue) and 2) in cells without AB co-repair (grey). Triplets are sorted on the *x*-axis based on the delta *Z*(Jaccard). **g**, Scatter plot combining both groups of **f**, so that each triplet is represented only once.

We reasoned that, in order for DSB pairs to physically interact upon DNA damage in a given cell, both sites should be simultaneously occupied by the repair machinery to gain affinity for one another. Genome-wide single-cell data offers a unique possibility to examine such coordinated protein binding, also between distant and inter-chromosomal loci. We applied the same bin-based coordination metric as in Figure 3: if two loci are both coordinately bound by repair protein across single cells, these loci show high correlation. Indeed, we find highly coordinated 53BP1 binding events between pairs of individual break sites, which strongly overlaps the enriched Hi-C damage-induced contacts (Fig. 4d). We systematically distinguish all contacting DSB pairs from those that do not contact and find coordinated 53BP1 binding exclusively at contacting DSB pairs (Fig. 4c bottom).

While this correlation analysis captures quantitative signals in individual genomic bins, it does not consider whether signal-containing bins are part of a consecutive sequence (as is the case in a repair domain). To know whether entire repair domains also behave in a coordinated manner, we turned to our MSR segment calls, which represent presence or absence of repair protein along the entire DSB region. Because these segment calls are binarized, we quantified coordination with the Jaccard index, which measures the similarity between two sets of observations. In this context, that is the overlap in presence of repair protein segment “A” and “B” among all cells. This analysis confirmed that contacting pairs show highly increased repair protein coordination, at the level of whole domains, in all DDR protein single-cell datasets (Extended Data Fig. 5c).

#### Coordination of single-cell repair protein binding is variable across cells

In the analyses described above for Figure 4, both the Hi-C contacts and repair coordination measure pairwise (two-way) events, meaning they are limited to two DSB sites at a time. Notably, the population-based Hi-C data indicates that various DSBs are in close physical proximity to more than one other DSB—some DSBs contact >15 other sites (Extended Data Fig. 5d). Consequently, it is currently unknown to what extent contacting sites form multi-way, or higher-order, DSB repair “hubs”. The simplest form of such a multi-way hub is a “triplet” of given DSB sites A, B, and C (Fig. 4e). In a triplet, Hi-C contacts are found between all three sites, forming pairwise combinations AB, AC, and BC. However, the question remains whether triplet ABC is formed within one cell, or represents mutually exclusive contacts that occur independently across cells.

Although scDamID and scChIC profiles do not measure spatial localization of DSBs, pairwise damage-specific contacts are prominently recapitulated. Hence, we reasoned we could test the premise of intercellular heterogeneity by quantifying repair coordination of DSB triplets. A triplet is considered “cooperative” if AB and BC are coordinated, and “competitive” if AB hinders BC. We find that triplets are decidedly more likely to show cooperative repair. This is indicated by higher coordination scores of BC given AB (blue) compared to BC given no AB (grey) (Fig. 4f). Still, some triplets are competitive, although these triplets show less coordination overall. Notably, cooperative and competitive behavior are highly anti-correlated, for all measured repair factors (Fig. 4g, Extended Data Fig. 5c). This suggests that, in most cells, DSBs are coordinately bound by repair protein within the triplet; only in very few cells will triplet sites be bound coordinately with another DSB (or remain unbound by repair protein altogether).

By distinguishing multi-way repair coordination within one cell from different combinations of pairwise coordination across cells, we find increased cooperative repair, in support of multi-way DSB repair hub formation.

### Repair protein foci cluster in multi-way chromatin hubs

To solidify our hypothesis that higher-order coordination of repair protein binding events suggests the presence of DSB repair contact hubs, we sought to prove that damaged sites form multi-way contacts on the genomic level. We addressed this gap by performing Tri-C experiments^57,58^ in damaged and undamaged conditions. Tri-C is a multi-way 3C approach, which enables identification of multiple ligation junctions in 3C concatemers. Since the order of fragments in these concatemers represents the 3D conformation of individual alleles, Tri-C gives insight into higher-order chromatin structures formed in single cells. By using capture oligonucleotide-mediated enrichment of regions of interest, Tri-C allows for analysis at high resolution and sensitivity. We designed 13 unique viewpoints (VPs) that each cover a single DSB site. The final DNA-sequencing reads are thus expected to contain a viewpoint fragment, and one or more proximal fragments, originating from single-allele chromatin conformations.

First, we used Tri-C to validate the presence of DSB-induced pairwise contacts between a given VP and other DSB sites. Both long-range and inter-chromosomal contacts are specifically formed in the damaged condition (Fig. 5a, Extended Data Fig. 6a-b). As intended, many of the Tri-C reads are composed of more than two contact fragments (Extended Data Fig. 6c). To analyze multi-way hub formation between DSB sites, we turned to the minimal multi-way hub case: a triplet of given DSB sites A, B and C (Fig. 4e). We defined the Tri-C experimental VP as site A, and performed in silico selection at site B (see Methods). Briefly: all reads containing site A are divided into two sets: one that contains only site A (negative selection), and one that contains both A and B (positive selection). We then visualized the contact profiles of the remaining proximal fragments. These fragments are specifically enriched at other DSBs (*i.e.*, sites C), thus forming three-way interactions on single alleles (Fig. 5b, Extended Data Fig. 6d). To quantify this observation, we calculated damage-specific contacts at site C, termed “Tri-C score”, for each possible triplet ABC (see Methods and Fig. 5b bottom).

**Fig. 5.**
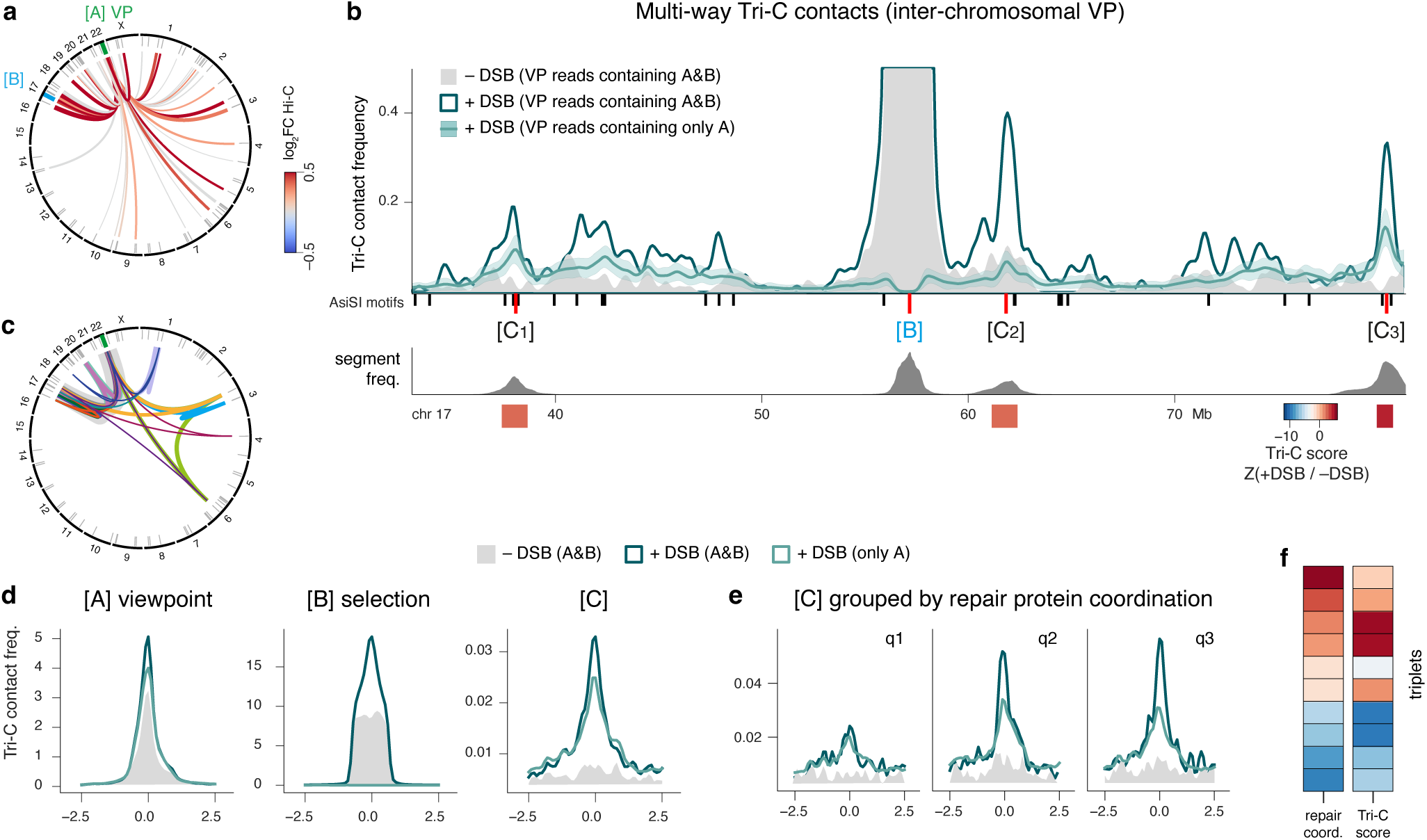
Damage-specific three-way contacts identified by Tri-C. **a**, Circos plot showing pairwise Tri-C contacts (lines) between the given VP and all other DSB top sites across the genome. The width of the lines represents the Tri-C contact frequency and the color represents the Hi-C fold-change between –DSB and +DSB conditions. Ticks indicate the DSB site locations. **b**, (top) Contact profile of positively selected reads that, aside from the VP (defined as site A), also have a fragment overlapping a second DSB site B. Conditions are –DSB (grey) and +DSB (teal). Conversely, contact profile of negatively selected reads, specifically lacking of fragment overlapping site B (light teal, mean ± SD). (bottom) Repair protein segment frequency of 53BP1. Below, repair domains (boxes) are used to quantify the Z-score at DSB sites [C] between positively and negatively selected profiles. **c**, Circos plot showing all three-way Tri-C contacts formed by the given VP, with a minimum Tri-C score of 0.5. Line width represents Tri-C score. Each color represents one three-way contact. **d**, Alignment plot of +DSB contact profiles as in b for all triplets of all 13 Tri-C viewpoints, where each time the VP was defined as site A, positive (teal) or negative (light teal) selection was done on site B, and the resulting contact profile is examined on site C. The grey contact profile is that of positive selection for site B in the –DSB condition. **e**, Same as the right panel of site C in c, but here stratified over 3 quantiles (q) that are based on increasing single-cell repair protein coordination. **f**, Heatmaps comparing single-cell repair protein coordination (left) to three-way Tri-C contacts (right). Coordination is measured as BC co-repair given AB co-repair. Tri-C contact is measured as Z-score of triplet site C. For this VP, all triplets are shown that contain a minimum of 30 reads at site C.

Using this measure, we could now visualize triplets formed with this VP across the genome (Fig. 5c). Our finding of three-way topologies is supported by all 13 Tri-C datasets (Fig. 5d). Finally, we aimed to integrate the multi-way Tri-C analysis with single-cell coordination of DSB repair protein. Triplets were grouped into three quantiles based on increasing repair coordination. Indeed, higher repair protein coordination correlated with increased Tri-C contact enrichment (Fig. 5e), even at the level of individual triplets (Fig. 5f). In sum, we present the first direct evidence of multi-way clustering in response to damage, at single-molecule resolution.

### Higher-order repair coordination in single cells increases with hub size

As described above, multi-way DSB repair hub formation is prominently observed within single cells and on single molecules. While Tri-C contacts are currently limited to predominantly three-way structures, genome-wide single-cell profiles allow for theoretically infinite combinations. Thus, we set out to evaluate higher-order repair coordination, using Hi-C as an independent and orthogonal measurement. Based on pairwise Hi-C data, we selected DSB sites that each form 3 or more proximal contacts (Extended Data Fig. 5d). Hierarchical clustering identified subsets of sites that frequently interact (Fig. 6a, Extended Data Fig. 7a; clusters indicating colored boxes). The clusters are remarkably mirrored by pairwise repair coordination across single cells (Fig. 6a). From this DSB Hi-C matrix, we identified a few hundred hubs of sites that have pairwise contacts between all participating loci. These hubs vary in size (*i.e.*, number of contacts), with some extremes consisting of up to 6 DSBs (Extended Data Fig. 7b).

**Fig. 6.**
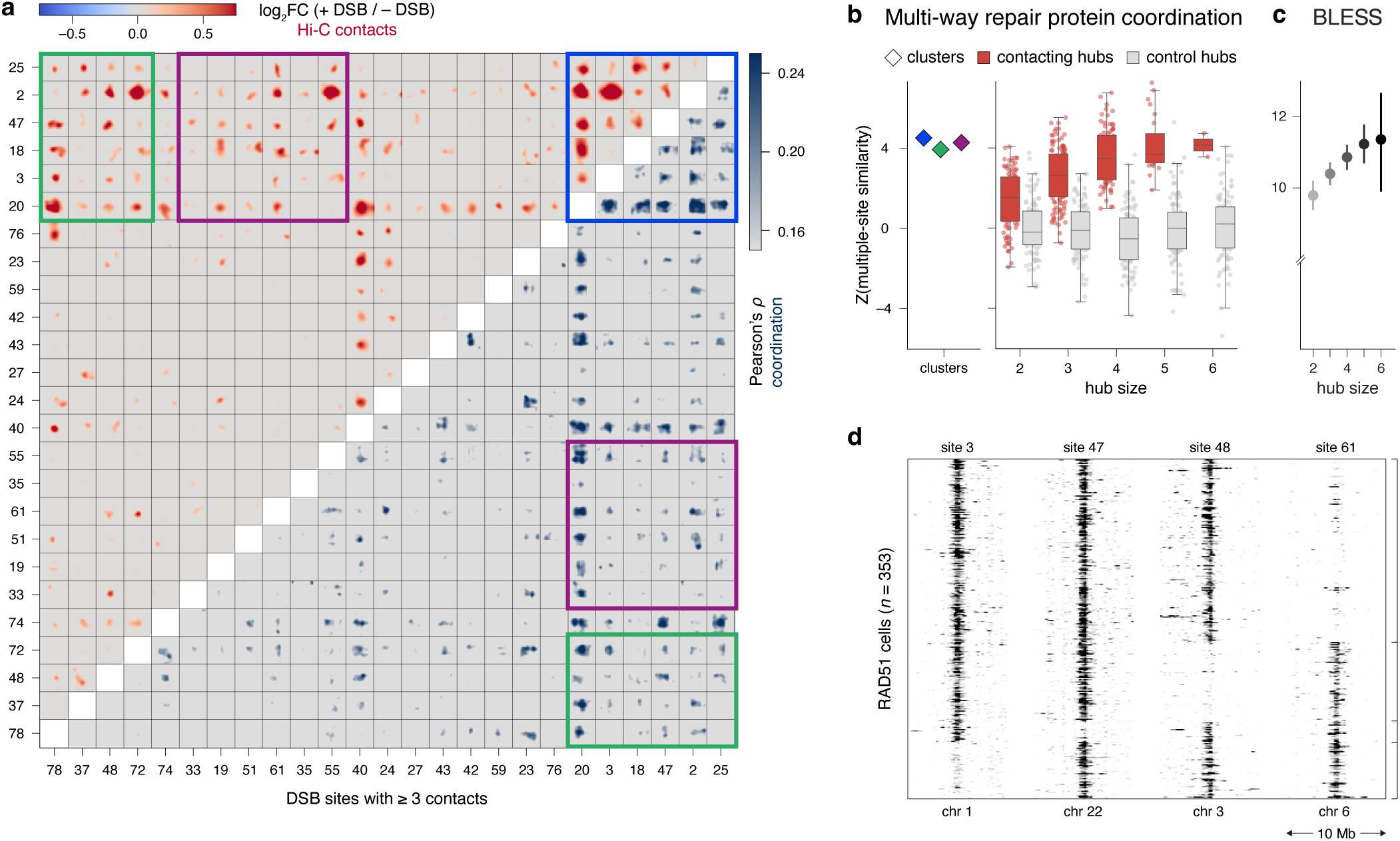
Higher-order coordination of repair protein binding in single cells. **a**, Clustered heatmap and hub annotation as in Extended Data 7a, displaying pairwise damage-specific Hi-C contacts (red, top left) and single-cell 53BP1 repair coordination (marine blue, bottom right), in a region of 1 Mb surrounding each AsiSI site. **b**, Higher-order repair protein coordination. Left: clusters corresponding to those identically colored in a. Right: Across all hub sizes, coordination measured for hubs in contact (red) and control hubs of the same size (gray). **c**, BLESS score (Aymard *et al*., 2017) measured across hub sizes. **d**, Heatmap showing single-cell Dam-RAD51 signal (RPKM) at 4 DSB sites in 10-Mb windows. Rows (*i.e.*, cells) are ordered according to different combinations of simultaneous RAD51 binding in each cell.

To directly test higher-order coordination in hubs, we applied a multiple-site similarity measure. For further explanation, see Methods and ^59,60^). First, we measured multi-way coordination in the large clusters identified from the Hi-C matrix. Indeed, these clusters showed very high coordination of repair protein binding (Fig. 6b, left). Next, in a systematic analysis, contacting hubs (red) showed considerably more multi-way repair coordination than control hubs of the same size (grey) that were randomly selected from all sites (Fig. 6b, right). Moreover, coordination consistently increased according to hub size, implying that cooperative repair (rather than multiple separate hubs) preferentially occurs within one cell. In support of this hypothesis, larger hubs show more frequent repair protein binding (Extended Data Fig. 7c). That is, a given DSB is bound in (many) more cells when that site is part of a larger hub. This is in line with our finding of cooperative versus competitive repair within triplets (Figure 4). In addition, larger hubs contain sites that are more frequently damaged, as shown by BLESS^61^, a method that detects genome-wide DSB distribution. Damage propensity can thus be interpreted as a source of variability in hub formation that is mimicked by repair protein recruitment and subsequent cooperative motion. Finally, we sought to explore variability in repair protein binding within hubs. A given hub of 4 DSBs (with pairwise Hi-C contacts between all) shows different combinations of simultaneous repair protein binding (Fig. 6d).

Our analysis thus shows evidence of cooperative and mutually exclusive higher-order DSB repair contacts, which are indistinguishable in Hi-C population data where all cases appear as pairwise interactions. Collectively, these data illustrate a range of repair coordination, with multi-way architectural repair hubs that may heterogeneously exist across single cells in various conformations.

## Conclusion

Above, we established the framework for sensitive, genome-wide, single-cell DNA repair factor profiling. Our experimental and computational approaches enable detection of repair protein enrichment at kilobase-scale resolution in thousands of individual cells. The sequence-specific DSB induction system allowed us to examine coordinated repair protein occupancy at known genomic positions. We show that repair proteins spread according to topological features but exhibit intercellular variability. Further, our analysis of multiple loci provides evidence of DSBs being simultaneously bound in single cells, information not currently attainable from spatial contact approaches. The observation of such coordinated repair protein binding is in line with recent studies suggesting that the physical properties of chromatin, along with compartmentalization and phase separation of repair proteins, facilitate the self-aggregation of certain damaged loci^56,62–65^.

The genomics toolbox now available enables study of other DNA damage and repair mechanisms, including stochastic systems and in vivo models. Stratification of single-cell DNA repair profiles according to an additional layer of information is amenable to many different questions of interest. We demonstrated the applicability by using live DNA content staining prior to index sorting to explore the cell cycle. A similar approach can be envisioned for any cellular observation that is compatible with FACS, e.g., antibody staining for cell type annotation, classification of apoptosis or other stress responses, and mitochondrial labeling. We foresee that (multimodal) single-cell measurements will disentangle the role of chromatin, transcription, and other factors involved in DNA repair at high resolution, with implications for our understanding of cellular response and fate after damage.

## Data availability

Newly generated sequencing data are publicly available on the NCBI Gene Expression Omnibus under accession code GSE229874. See Methods for accession numbers of publicly available sequencing data analyzed in this manuscript. Imaging data will be made publicly available on Mendeley Data.

## Code availability

Key scripts will be made available on GitHub / Zenodo. See www.github.com/KindLab/scRepair.

## Acknowledgments

We thank members of the Kind and Legube groups and Eva Brinkman for feedback throughout the project. Specifically, we acknowledge Koos Rooijers for developing computational methods and performing bioinformatic analyses. Purified pA-MNase and m6A-Tracer were respective gifts from Alexander van Oudenaarden and Bas van Steensel. We thank Reinier van der Linden at the Hubrecht FACS Facility and Anko de Graaff at the Hubrecht Imaging Center for technical support and training. High-throughput sequencing and computational resources were provided by the Utrecht Sequencing Facility (USEQ) and High-Performance Computing Facility. USEQ is subsidized by the University Medical Center Utrecht and the Netherlands X-omics Initiative (NWO project 184.034.019). SPRITE experiments performed in the Jachowicz lab were supported by OEAW core funds. AMO acknowledges funding from the Max Planck Society. The Oncode Institute is supported by the KWF Dutch Cancer Society. This work was further funded by: Dutch Research Council (NWO-ALW) Vidi grant 161.339 (JK), Fulbright Commission the Netherlands promovendus grant (KLdL), European Molecular Biology Organization grant 9178 (KLdL).

## Author contributions

Conceptualization: KLdL and JK. Methodology, Investigation, Data curation & Validation: KLdL. MAK contributed Tri-C experiments, advised by AMO. SSdV performed revision experiments with KLdL. AGM and LSP respectively contributed SPRITE experiments and data processing, advised by JWJ. Formal analysis & Visualization: KLdL and PMJR. Software: PMJR. Project administration: JK. Resources: GL, JWJ, AMO, and JK. Funding acquisition: KLdL, JWJ, AMO, and JK. Writing – original draft: KLdL with contributions from PMJR and JK. Writing – review & editing: all authors.

## Competing interests

The authors declare no relevant competing interests.

---

**Extended Data Fig. 1.**
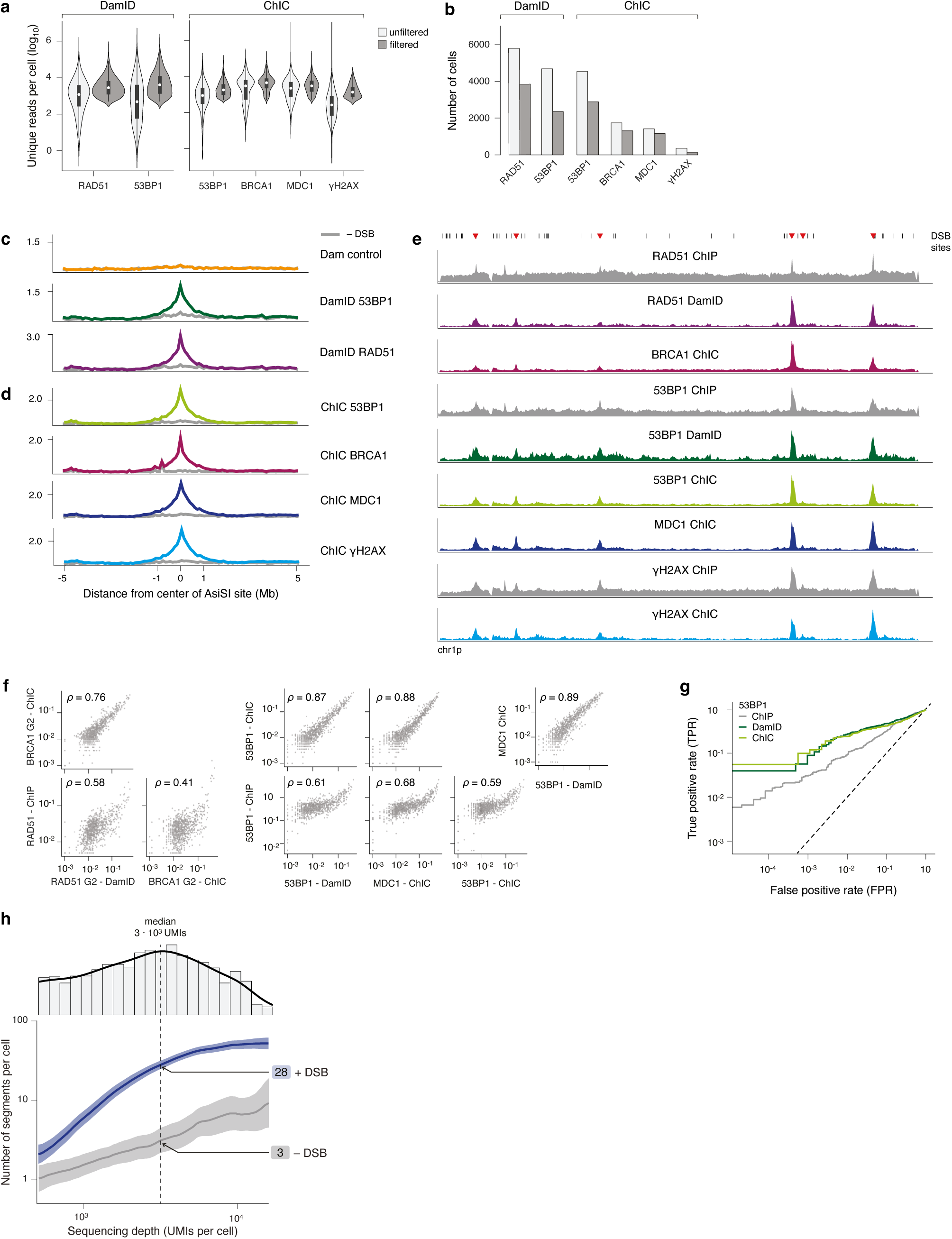
Technical validation of single-cell DamID and ChIC repair protein data. **a**, Distribution of sequencing depth per protein target per method, unfiltered and filtered (retaining samples passing threshold). **b**, Number of cells per protein target per method, unfiltered and filtered. **c**, DamID alignment plots (averaged across single cells) over top AsiSI sites, in +DSB (colored) and –DSB (gray) conditions. **d**, ChIC alignment plots (averaged across 100-cell samples) as in **c**. **e**, Plots of chromosome 1p as in Fig. 1b, incl. external ChIP-seq data. **f**, Scatter plots of ChIP RPKM versus DamID and ChIC repair protein frequency, per protein and relevant cell cycle phase. Each dot is an AsiSI site. Correlation values indicate Spearman’s *rho*. **g**, Receiver-Operator Characteristic (ROC) curves showing true positive rate (TPR) versus false positive rate (FPR) for 53BP1 ChIP, ChIC and DamID, testing presence of a called repair protein segment and whether it falls on an AsiSI motif. Dashed line: *x* = *y*. **h**, Line graphs showing the number of enriched segments as a function of sequencing depth in both DSB induction conditions. Bold line indicates mean, shaded area indicates 95% CI. Histogram of sequencing depth is annotated on top, dashed vertical line indicating the median.

**Extended Data Fig. 2.**
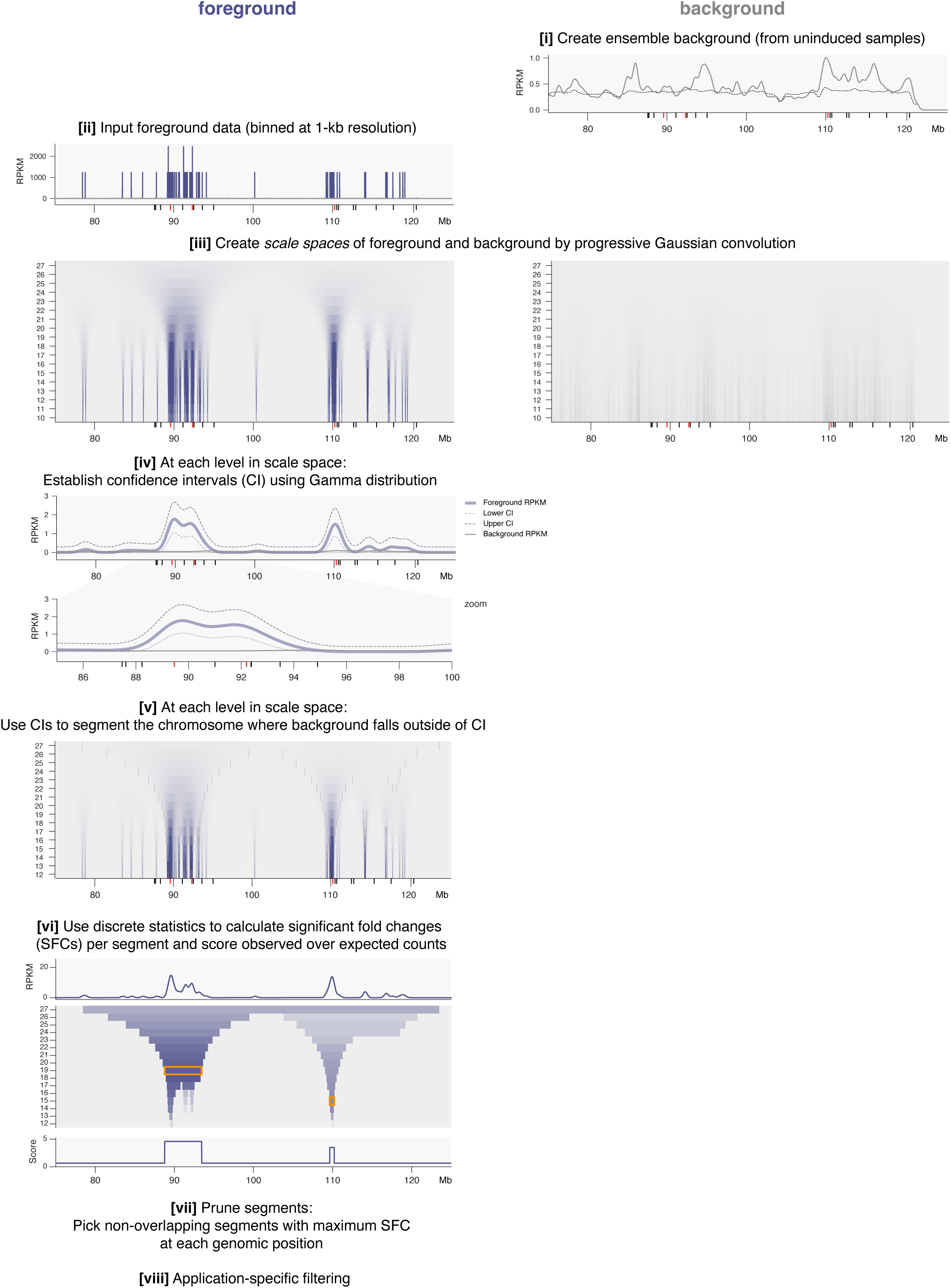
Computational model for unbiased detection of enriched genomic segments.

**Extended Data Fig. 3.**
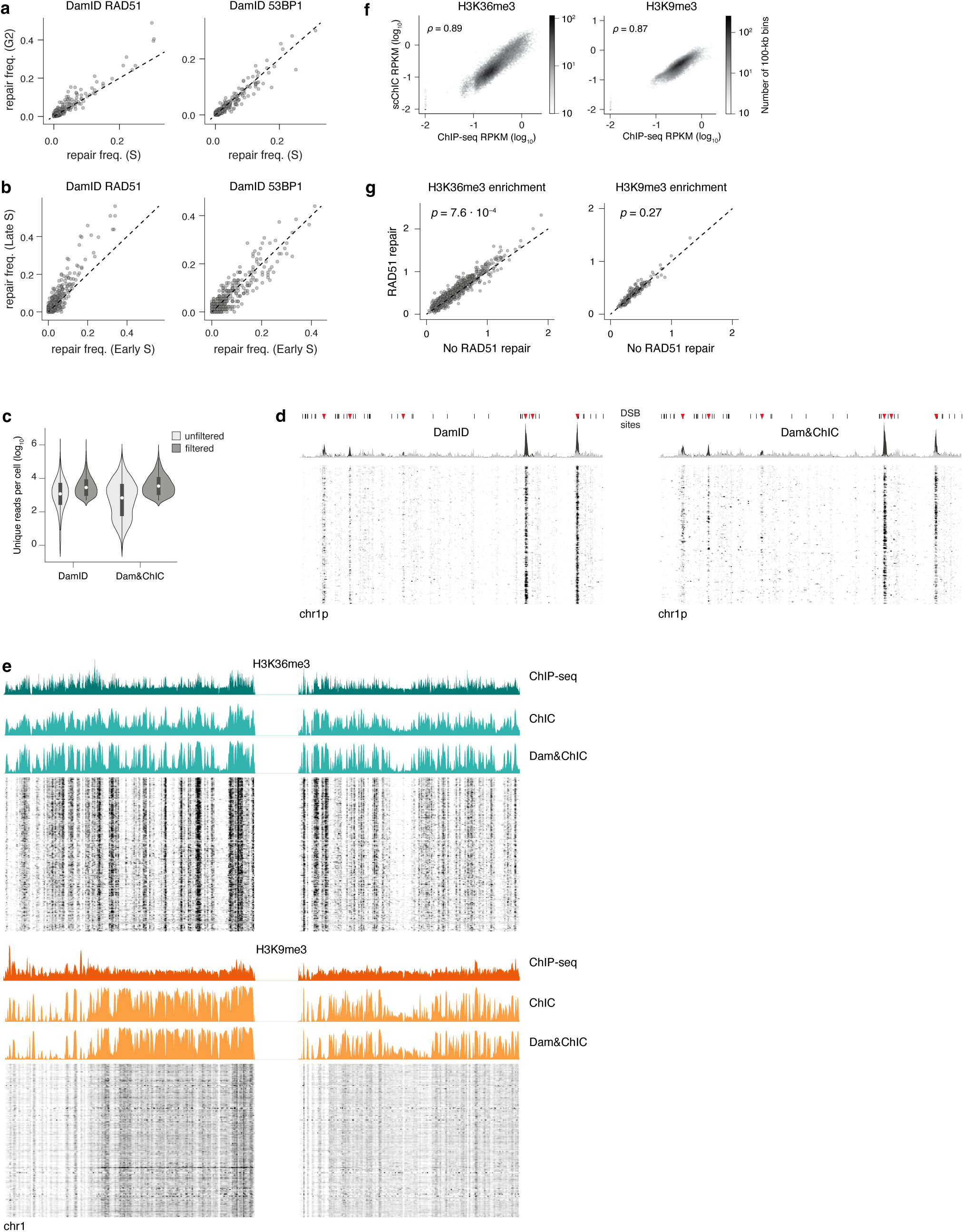
Validation and characterization of pathway-specific repair protein frequencies. **a**, Scatter plots of RAD51 and 53BP1 DamID repair protein frequency in G2 vs S phase. Each dot is an AsiSI cluster. **b**, Scatter plots as in **a**, showing Late vs Early S phase. **c**, Sequencing depth (UMIs per cell) for DamID or Dam&ChIC samples. **d**, DamID RAD51 signal on chromosome 1p, generated with protocols as in **c**. **e**, Signal on chromosome 1, for H3K36me3 (top) and H3K9me3 (bottom). Line plots show ChIP-seq (RPKM) and single-cell aggregate ChIC (contact frequency). Heatmaps show scChIC (RPKM) of 1000 cells. Aggregate ChIC signal is split on sample preparation by ChIC or Dam&ChIC. **f**, Hexbin density plots showing the bin-based correlations between ChIP-seq and scChIC sequencing depth. Hexbins are colored by the number of 100-kb genomic bins. **g**, Genome-wide quantification of H3K36me3 (left) or H3K9me3 (right) signal based on presence or absence of RAD51 enrichment. For each AsiSI site, scatter plots show the mean hPTM ChIC signal in cells that are repaired by RAD51 (*y*) or not (*x*). Statistical significance was calculated by one-sample KS test.

**Extended Data Fig. 4.**
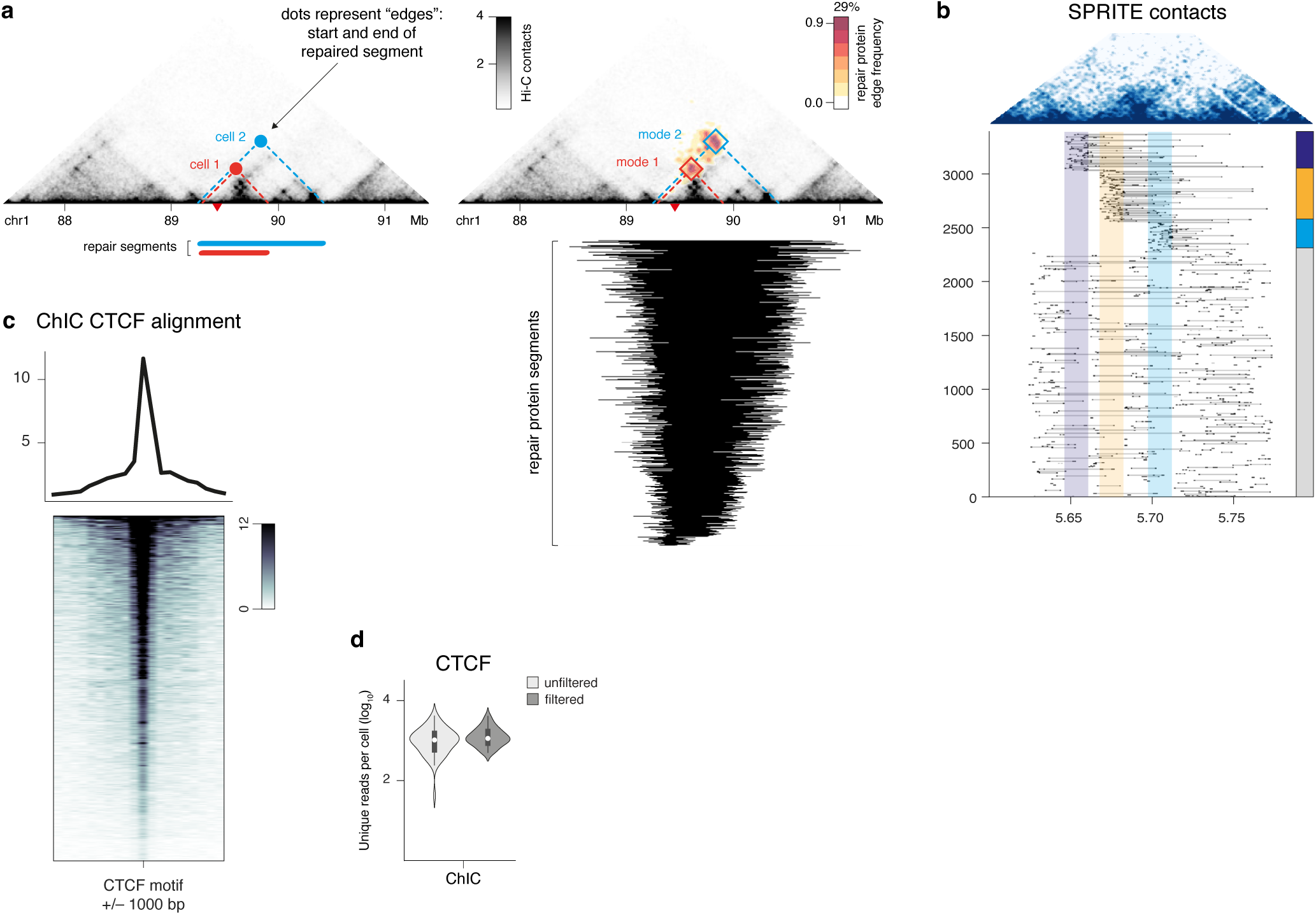
Intercellular variability visualized by projection of repair protein segment edges onto Hi-C contact profiles. **a**, Overlay of repair protein segment edges onto the Hi-C contact matrix for regions harboring a DSB site. Left: cartoon indicates repair protein segments of different sizes, originating from two individual cells. Right: Each repair data point projected onto the Hi-C map represents the genomic coordinates of a segment start and end. Repair protein signal is colored by the frequencies with which those edges are observed across single cells. Repair protein segments (as called on the linear genome) are annotated underneath, sorted on segment size (large to small, top to bottom). **b**, Pairwise SPRITE contact map (top) of the site shown in **3a**. Individual SPRITE clusters (rows) are plotted underneath, aggregated by the three repair protein spreading scenarios as in **3b**. **c**, ChIC CTCF alignment plots (averaged across single cells) over CTCF motifs +/– 1000 bp. **d**, Distribution of ChIC CTCF sequencing depth, unfiltered and filtered (retaining samples passing threshold).

**Extended Data Fig. 5.**
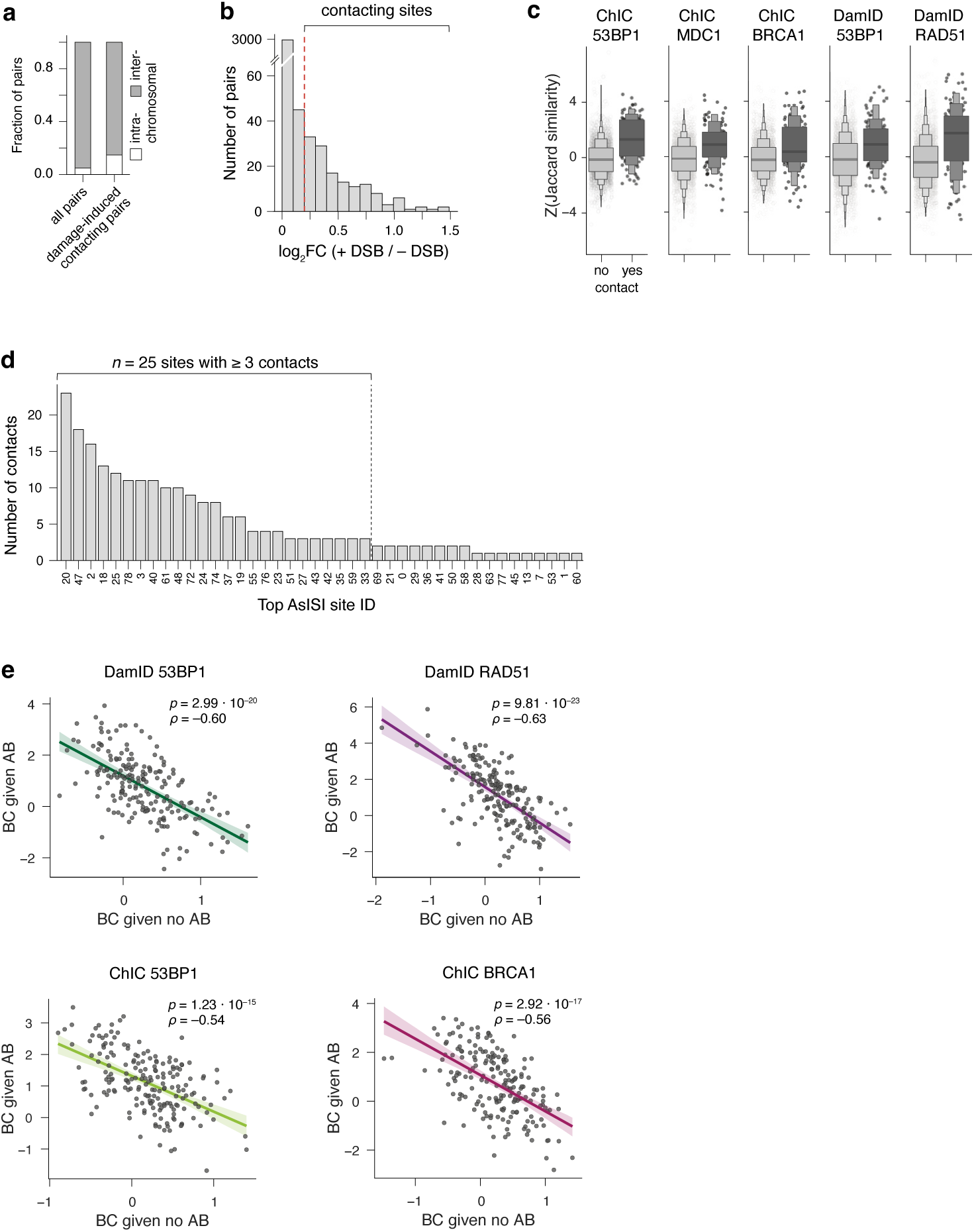
Characterization of pairwise contacts and associated repair features. **a,** Quantification of (damage-specific) contacting pairs as intra- or inter-chromosomal. **b**, Histogram showing the number of DSB pairs across Hi-C contact scores (log_2_FC (+DSB / –DSB)). **c**, Pairwise single-cell coordination of binarized repair enrichment (MSR calls). Coordination is measured with the normalized Jaccard index, for pairs categorized as in **4d**. **d**, Bar plot showing number of contacts formed per site. **e**, Scatter plot indicating pairwise similarities of BC for both cases of AB. Same as in **4g** for the indicated repair proteins.

**Extended Data Fig. 6.**
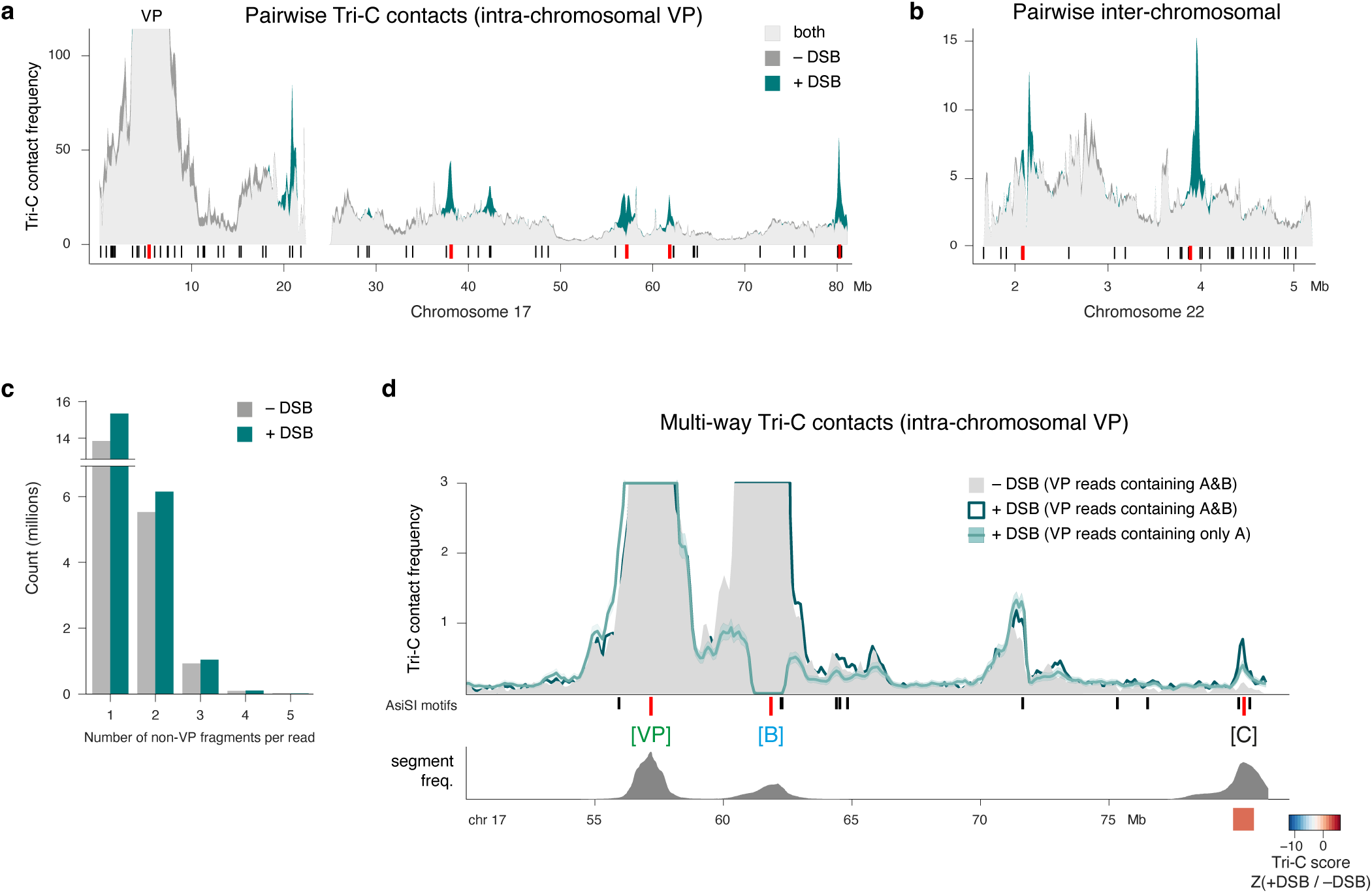
Characterization of Tri-C contacts. **a-b**, Pairwise Tri-C contact profile in –DSB (grey) and +DSB (teal) conditions. Ticks underneath indicate location of DSB sites with previously annotated top sites in red. Intra-chromosomal contacts on chr 17 in **a**, and inter-chromosomal contacts on chr 22 of the same VP in **b**. **c**, Histogram showing the distribution of reads according to the number of non-viewpoint fragments per read. **d**, Multi-way Tri-C contact profile as in **5b**, but showing an intra-chromosomal VP.

**Extended Data Fig. 7.**
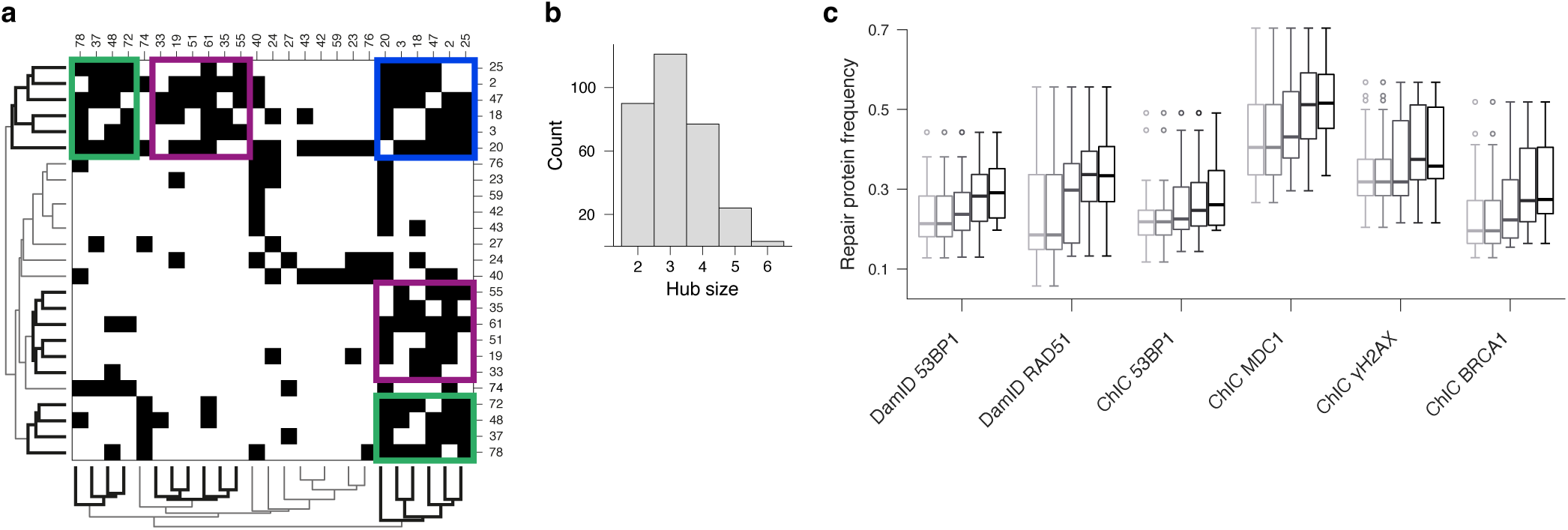
Characterization of higher-order contacts and associated repair features. **a**, Heatmap of all sites forming three or more contacts, clustered based on binarized contact score. Colored rectangles indicate hubs of sites that display increased multi-way contacts between them. **b**, Histogram showing the number of formed hubs containing 2-6 sites. **c**, Repair protein frequency measured across hub sizes, for all datasets.

## Methods

### Experimental methods

#### DamID construct design

Shield1-inducible Dam lines were created by cloning into the multiple cloning site (MCS) downstream of the double degron (DD) of ProteoTuner vector pPTuner IRES3 (PT4040-5). This generates an in-frame construct of DD-POI-V5-Dam. POI 53BP1 contains the minimal focus-forming region (amino acids 1221-1711). POI RAD51 contains the full endogenous protein (amino acids 1-339).

#### Generation of cell lines

Stable, clonal cell lines containing Dam-POI constructs were established by transfection and antibiotic selection. DIvA cells were grown in 24-well plates and transfected with 500 ng DamID plasmid and 1.5 µL Lipofectamine 2000 per well. Each well was passaged to a 15-cm dish and subjected to antibiotic resistance selection with 500 µg/mL G418 (Gibco) for 10 days (at complete death of untransfected control dishes). Monoclonal cell populations were hand-picked, expanded, and characterized by performing bulk DamID. Dam methylation levels were checked by evaluating methylation-specific amplification on agarose gel (as previously described in ^14,66^) and on-target methylation signal was evaluated with high-throughput sequencing. One clone per construct with the highest signal-to-noise ratio (SNR) was chosen for single-cell experiments.

#### Cell culture and experimental treatment conditions

Cell lines were grown in DMEM containing high glucose, GlutaMAX supplement, and sodium pyruvate (Gibco) supplemented with 10% fetal bovine serum (FBS; Sigma-Aldrich) and 1X Penicillin Streptomycin (Gibco), at 37 °C with 5% CO_2_. For maintenance, cells were split 1/10 every 3 days and routinely tested for *Mycoplasma*. For experimental procedures, cells were plated the day prior, to achieve 60-70% confluency during induction treatment. Dam-POIs were stabilized by addition of 500 nM Shield1 ligand (AOBIOUS) for 4 h (Dam-53BP1) or 8 h (Dam-RAD51). Nuclear translocation of AsiSI-ER was simultaneously induced by addition of 300 nM 4-hydroxytamoxifen (4OHT; Sigma, dissolved in ethanol to 13 mM). Cells were harvested by washing twice with PBS0 followed by trypsinization with 1X TrypLE (Gibco), inactivation with DMEM, pipetting to yield a single-cell suspension, and centrifugation at 300 *g*. For FACS, cell pellets were resuspended in growth medium containing 10 µg/mL Hoechst 34580 (Invitrogen) per 1x10^6^ cells and incubated for 45-60 minutes at 37 °C, for live DNA content staining. Prior to sorting, cell suspension was passed through a 20-µm mesh. For bulk DamID, genomic DNA was isolated from cell pellets using commercial reagents (e.g., Promega Wizard). During each genomics experiment, cells were concurrently plated on glass and treated to verify proper induction and cleavage activity of AsiSI-ER by immunofluorescent staining and imaging.

#### Immunofluorescent staining

Cells were grown as described above, with the exception of plating on glass coverslips the day prior to experimental treatment. At the end of induction, cells were washed twice with PBS and chemically crosslinked with fresh formaldehyde solution (2% in PBS) for 10 minutes at RT, then permeabilized (with 0.5% IGEPAL® CA-630 in PBS) for 20 minutes and blocked (with 1% bovine serum albumin (BSA) in PBS) for 30 minutes. All antibody incubations were performed in final 1% BSA in PBS followed by three PBS washes at RT. Incubation with primary antibody against the endogenous protein as well as purified ^m6^A-Tracer protein (recognizing methylated DNA) was performed at 4 °C for 16 hours (overnight), followed by anti-GFP (against ^m6^A-Tracer protein) incubation at RT for 1 hour, and secondary antibody incubations at RT for 1 hour. The final PBS wash was simultaneously an incubation with DAPI (Invitrogen) at 0.5 μg/mL for 2 min, followed by a wash in MilliQ and sample mounting on glass slides.

Primary antibodies: anti-53BP1 Santa Cruz (rabbit) at 1/500, anti-γH2AX BioLegend (mouse) at 1/1000, anti-MDC1 Bethyl Laboratories (rabbit) at 1/500, anti-GFP Aves GFP-1020 (chicken) at 1/1000.

Secondary antibodies: all AlexaFluor at 1/500. Anti-chicken 488, anti-mouse 555 or 647, anti-rabbit 555 or 647.

#### Confocal imaging

Imaging (12-bit) was performed on an inverted scanning confocal microscope (Leica TCS SP8) with a HC PL APO CS3 63X (NA 1.40) oil-immersion objective and HyD detectors. Pinhole was set to 1 Airy Unit. Scanning zoom was set to 4-5X at a speed of 400 Hz. Full-nucleus images were acquired as *Z*-stacks at 0.2-μm intervals. Multi-color images were acquired sequentially (by frame).

#### Image analysis

Images were processed in Imaris 9.3 (Bitplane) by baseline subtraction and background correction with a 3x3(x3) median filter. Colocalization was calculated by Manders’ coefficients M1 and M2 between channel 2 (DamID ^m6^A-Tracer) and channel 3 (endogenous DSB repair marker). Pixels were retained that contained signal (intensity >10) in channel 1 (DAPI). Pixel intensity thresholds for the colocalization analysis were determined using Costes’ method with default settings.

#### FACS

FACS was performed on BD FACSJazz or BD FACSInflux Cell Sorter instruments with BD Sortware. Index information was recorded for all sorts. Single cells were gated on forward and side scatters, trigger pulse width, and Hoechst cell cycle profiles. One cell per well was sorted into 384-well hard-shell plates containing 5 μL of filtered mineral oil and protocol-specific reagent.

#### DamID and derivative methods

##### High-throughput sequencing

Libraries were sequenced on the Illumina NextSeq 500 (75-bp single-end reads) or NextSeq 2000 (100-bp paired-end reads) platform.

##### Bulk DamID

DamID on populations was performed as previously described^66^. Briefly, genomic DNA (gDNA) was isolated from cell pellets, digested with DpnI to enrich for Dam-methylated GATCs, and ligated to universal (not barcoded) double-stranded DamID adapter molecules. Methylation-specific PCR was performed with barcoded primer (unique per sample). Samples were pooled per clone, further processed to construct Illumina-compatible libraries, and sequenced to approximately 10 M raw reads per sample.

##### Automated liquid handling

Liquid reagent dispensing steps for single-cell protocols in microwell plates were performed on a Nanodrop II robot (Innovadyne Technologies / BioNex). Addition of barcoded adapters was done with a mosquito LV (SPT Labtech).

##### Single-cell DamID2

DamID on single cells was performed as previously described in detail^67^. Briefly, after FACS, cells were lysed and treated with proteinase K, after which methylated GATCs resulting from Dam enzyme activity were specifically digested with DpnI. Double-stranded adapters containing cell-specific barcodes and a T7 promoter were ligated to the blunt (DpnI-cleaved) DNA ends. Cells with non-overlapping barcodes were pooled together to undergo *in vitro* transcription (IVT), amplifying the genomic DamID-specific product in a linear manner. Library preparation was then performed on the amplified RNA, to generate molecules compatible with Illumina sequencing.

##### Single-cell Dam&T-seq

One single-cell DamID experiment was performed using the combinatorial scDam&T-seq approach capturing DamID and transcriptome ^15^, as previously described in detail ^67^, with the exception that all volumes were halved (to reduce costs). Briefly, after FACS, cells were lysed, followed by reverse transcription and second-strand synthesis in order to convert cellular mRNA into cDNA. Subsequent steps were followed according to the scDamID2 protocol.

##### (Single-cell) ChIC

ChIC was performed as described in detail in ^10^, with adaptations as follows. After experimental treatment of cell cultures as described above, nuclei were isolated and permeabilized, incubated with primary antibody, then incubated with pA-MNase (Protein A IgG-binding domain fused to micrococcal nuclease, for antibody-specific binding) and Hoechst (for DNA content staining). If the primary antibody was raised in mouse, nuclei were incubated with secondary antibody (rabbit anti-mouse) before incubation with pA-MNase. After FACS, proximity-based cleaving by pA-MNase was activated (for exactly 30 min on ice), followed by inactivation and proteinase K treatment. MNase-cleaved ends were then blunted and phosphorylated, and double-stranded adapters were ligated. For one experiment, A-tailing was performed after end repair of MNase-cleaved ends, followed by ligation of T-tailed adapters. DNA molecules were then further processed for sequencing as in scDamID2.

Antibodies: anti-53BP1 Santa Cruz [H-300] sc-22760 (rabbit) at 1/500, anti-γH2AX BioLegend [2F3] 61340x (mouse) at 1/500, anti-MDC1 Bethyl Laboratories A300-053A (rabbit) at 1/500, anti-BRCA1 Santa Cruz [D-9] sc-6954 (mouse) at 1/500, anti-H3K9me3 Abcam ab8898 (rabbit) at 1/1000, anti-H3K36me3 Active Motif 6110x (rabbit) at 1/500, anti-mouse IgG Abcam ab6709 (rabbit) at 1/500, anti-CTCF Merck 07-729 (rabbit) at 1/200.

##### Single-cell Dam&ChIC

Combinatorial profiling of DamID and ChIC was performed as described in ^11^, with adaptations as follows. The same procedures of nuclei isolation, antibody treatment, FACS, and molecular preparation were followed as for scChIC. After end repair of MNase-cleaved ends, the scDamID procedure was followed, namely DpnI digestion to enrich for Dam-methylated GATCs, adapter ligation, and subsequent library preparation steps.

##### Tri-C

Measurement of multi-way contacts with Tri-C was performed following previously published protocols ^58^, in 2 biological replicates per experimental condition and 4 technical replicates per biological replicate. Briefly, cells were collected in culture medium in batches of 15 M cells per technical replicate and cross-linked for 10 minutes with 2% formaldehyde (ThermoFischer, 28908). To prepare 3C libraries, aliquots of cells were split equally into 3 reactions and digested with NlaIII enzyme (NEB, R0125L). Then, a proximity ligation was performed and ligated chromatin was extracted with Phenol-Chloroform method. The separate digest reactions were combined. 8 µg of 3C library per technical replicate was sheared with Covaris S220 Focused-Ultrasonicator to the mean size of 450 bp (time: 55 s, duty factor: 10%, peak incident power: 140 W, cycles per burst: 200). In order to exclude fragments shorter than 300 bp, which are unlikely to contain more than one ligation junction, the samples were size-selected with 0.7x of Mag-Bind TotalPure NGS beads (Omega Bio-Tek, M1378-01). Samples were indexed in duplicate in order to increase sample complexity using NEBNext Ultra II DNA Library Prep Kit for Illumina (NEB, E7645S), and Herculase II polymerase (Agilent, 600677) for sample amplification.

Indexed libraries were enriched for the viewpoints of interests in a double-capture procedure. Probes used for capture were designed with python-based oligo tool (https://oligo.readthedocs.io/en/latest/). The 120 nt long, 5’-biotinylated, ssDNA probes were ordered as a multiplexed panel of oligos (IDT, xGen™ Custom Hybridization Capture Panels), and used at 2.9 nM concentration. The enrichment was performed using the KAPA Hyper Capture Reagent Kit (Roche, 9075828001). A total of 12 µg of indexed sample per biological replicate was used as input for the first capture. Captured DNA was pulled down with M-270 Streptavidin Dynabeads (Invitrogen, 65305), washed, and PCR amplified. All recovered material was used as input for the second capture. The quality of final samples was assessed by fragment analyzer and samples were sequenced on an Illumina platform with 300 cycles paired-end reads.

##### SPRITE

SPRITE was performed as described in ^68^ with minor modifications. U2OS cells were crosslinked with DSG/1%PFA, followed by permeabilization and nuclear extraction according to the protocol. Next, nuclei were digested with a mix of restriction enzymes HpyCH4V and AluI for 16 hours, washed in PBStb buffer and sonicated using a Covaris E220. From this step onwards, the standard SPRITE protocol^68^ was followed. Sequencing was performed using the Illumina NovaSeq X platform (10B chemistry), with 500 million reads obtained.

### Computational methods

#### Raw data processing

Data generated by the scDamID and scDam&T-seq protocols was largely processed with the workflow and scripts described in ^67^ (see also www.github.com/KindLab/scDamAndTools). The procedure is described below, in brief. For detailed parameters and exact software versions, see www.github.com/KindLab/scRepair.

##### DamID and Dam&T samples

DamID data contains reads that result from DpnI-restriction activity. Briefly, raw sequencing data was processed by removing reads that contained contaminants (resulting from library preparation), using `cutadapt` (v2.0). Samples were then demultiplexed (using an in-house python script), and for every single sample one (or two, in the case of paired-end data) files were obtained. Demultiplexed data were aligned using `hisat2` (v2.1.0) to human reference genome hg19/GRCh37, supplemented with the ERCC spike-in sequences. Since DpnI cleaves GĂTC, we prefixed each read *in silico* with a ‘GA’ dimer to improve alignment rates. Aligned reads were counted per genomic GATC position and a vector of counts per chromosome was stored. For DamID, reads not aligning at GATC positions were discarded. Reads were then binned in either 1-kb or 100-kb bins for further analysis and plotting.

##### ChIC and Dam&ChIC samples

Contaminant removal, demultiplexing and alignment was done as for DamID samples. Reads were counted per GATC position (for both ChIC-only and Dam&ChIC samples), but reads not aligning on GATC position, as well as unaligned reads were stored in an auxiliary BAM file. These non-GATC reads were processed by removal of the (*in silico* prefixed) ‘GA’ dimer, and realignment to the human reference genome. Realigned non-GATC reads were then counted UMI-unique per genomic position and counts were binned in either 1-kb or 100-kb bins for further analysis and plotting.

##### SPRITE samples

Raw sequencing data were analysed based on ^45^ using the available pipeline (https://github.com/GuttmanLab/sprite-pipeline) and reference genome hg38. The pipeline was modified to enable local alignment using the “`--local`” flag of bowtie2. The barcode sequence for creating the SPRITE clusters was `[Y|SPACER|ODD|SPACER|EVEN|SPACER|ODD]`.

##### Tri-C samples

Tri-C data were processed using the capcruncher pipeline ^58^ (v.0.3.11) in tiled mode.

#### RPKMs and scaling factors

RPKMs were only used for inspection of raw read densities and were calculated using the common definition: 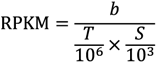 where *b* is the number of reads per bin, *T* is the sample total and *S* is the size of bins (in kb).

The nature of DSB binding patterns make the signal generally “inherently peaky” (high variance in read density along linear chromosomes). However, the number of DSBs and the ratio of signal (signal from DSB loci) and background (signal from inter-DSB loci) varies substantially between samples (*i.e.*, SNR, signal-to-noise ratios). Naively scaling the read density as with RPKMs led to false enrichments/depletions, e.g., comparing a high SNR sample to an uninduced (no DSBs) background would register depletions across most inter-DSB regions. To overcome the inherent variation in SNR between the single-cell samples we employed the “scaling factor” normalization from PoissonSeq^69^, section 3.2, which weakens the assumptions implicit of RPKM scaling by assuming only that a fraction (here: 50%) of bins are not differentially enriched between two conditions.

#### Enriched segment calling

To identify regions of significant enrichment and depletion we adapted the workflow of multi-scale representation of genomic signals as described in ^19^. A diagram of the workflow can be found in Extended Data Figure 2. Each induced sample is compared to a suitable background. For DamID, this is an average of single-cell samples that are not induced with 4OHT (one background per clone). For ChIC, this is an average of single-cell samples that are not induced with 4OHT (one per antibody target). Starting from 1-kb bins, both foreground and background signals are convolved with a Gaussian kernel which standard deviation increases with *√2* at each level, up to a standard deviation of 10 Mb. This creates the scale-space. Each level of the scale-space is segmented along the linear chromosome by comparing where background signal is either above or below the foreground signal (in practice a small confidence interval is built around the foreground signal to deal with zero data in the foreground signal and numerical rounding issues after Gaussian convolution, raising the possibility that background signal is *within* the confidence interval of the foreground signal; those segments are ignored in further steps). Each segment, where background signal is either below or above the foreground signal, at each level in the scale space is then tested for significant enrichment or depletion, respectively. We use the Gamma distribution to create a confidence interval around the background, using the observed signal density and the scaling factor (see above) of the foreground versus the background signal, and a *P*-value of *10^−9^*. A segment is considered significantly enriched or depleted when the foreground signal is either above or below this confidence interval. Similar to the original MSR, we calculate the “significant fold change” (SFC) as the observed foreground signal over the confidence interval boundary. In addition, we also record the true observed fold change of observed versus expected densities. Finally, we prune the enriched/depleted segments across all levels of the scale-space by selecting the level that yields the highest (absolute) SFC score.

##### Post-hoc filtering

We focus on “high-fidelity” enrichments in downstream analyses, satisfying the following conditions: observed logFC ≥1.25, size ≤10 Mb, segments encompass at least 10 UMI-unique reads in foreground sample.

#### Normalization of Hoechst measurements across batches

A Gaussian mixture model with 2 or 3 components, depending on fit, was applied to Hoechst intensity values, after which G1 and G2 peaks were assigned. Script is available on GitHub / Zenodo.

#### Defining top AsiSI sites

Top AsiSI site annotations were taken from ^33^.

#### External data processing

##### ChIP-seq

Raw ChIP-seq data was obtained from GSE48423, GSE97589 and E-MTAB-5817. Reads were aligned using HISAT2 (same parameters as our DamID and ChIC samples). Aligned reads with MAPQ ≥10 were counted in genomic bins of either 1 kb or 100 kb and used for downstream analyses.

##### Repli-seq

Repli-seq data was obtained from the 4DN project, SRP126407 and SRP197558. Samples used:

4DNFI6KIPWXQ - 4DNFID41JKT6 - 4DNFIFEZTAI1 - 4DNFIZDPE9T6.

SRR6363337 - SRR6363338 - SRR6363341 - SRR6363342 - SRR6363345 - SRR6363348 - SRR6363350 - SRR9040713 - SRR9040714.

Sequencing data was aligned using HISAT2 using the same parameters as DamID and ChIC samples. Reads were counted in bins of 1 kb. Using sample annotations of early and late replication (and for one sample set, mid-replication) we used glmPCA^56^ to obtain a principal component per chromosome indicative of replication timing in U2OS cells.

##### Hi-C

Files in `.hic` format were obtained from E-MTAB-8851. Samples used:

HiC_mOHT_rep1 - HiC_mOHT_rep2 - HiC_pOHT_rep1 - HiC_pOHT_rep2.

The observed over expected Hi-C matrix presented in Figure 3F was calculated by dividing the normalized Hi-C matrix (binned at 25-kb resolution) by the average intrachrotmosomal contact probability between increasingly distant pairs of loci (*i.e.*, distance-dependent contact decay) using `cooltools.lib.numutils.observed_over_expected`.

For (Extended Data) Figure 4, the normalized Hi-C matrices (binned at 25-kb resolution) were smoothed using a Gaussian kernel (*σ* = 200 kb). The fold change was calculated (by dividing the smoothed induced matrix over the uninduced) and subsequently log_2_-transformed, after which insignificant logFCs were masked from the interaction maps.

#### hPTMs in cells with and without RAD51 repaired sites

The analysis presented Figure 2E and S2F, that demonstrates enriched H3K36me3 on AsiSI sites in cells where sites are repaired by RAD51, was performed as follows. Per site, cells were stratified by the presence or absence of an MSR segment call. Only sites that overlap with an MSR segment in at least 5 cells were included. In the resulting two groups of cells (*i.e.*, with and without repair), the mean enrichment of either H3K36me3 or H3K9me3 ChIC signal was calculated in the consecutive non-overlapping 100-kb bin wherein the geometric mean of the AsiSI cluster was located.

#### MSR segment edge peak calling

For systematic comparison between MSR segment edge frequency and Hi-C insulation score (computed using `cooltools.insulation` on 25-kb binned normalized Hi-C matrix) presented in Figure 3C-D, segment edge peaks were identified as follows. The segment edge frequency was calculated by averaging the binary MSR calls over the single cells in 25-kb bins. The edge frequency vector was smoothed by fitting a Gaussian kernel (SD = 2.5 × 10^4^). Segment edge frequency peaks were subsequently detected by `scipy.signal.find_peaks` (with prominence = *P*_i_ where the *i*-th *P*ercentile is 97.5). Hi-C insulation score and segment edge frequency at called peak positions were used.

#### SPRITE analysis

We extracted SPRITE clusters from the aligned data using a modified version of get_sprite_contacts.py (https:// github.com/GuttmanLab/sprite-pipeline). Only sprite clusters containing between 2-10000 fragments were included for visualization. In Extended Data Fig. 4b all SPRITE clusters overlapping the locus are plotted and clustered according to overlap with repair protein segment edge peaks highlighted in Fig. 3a.

#### Identification of pairs of contacting repair sites

For the identification of contacting pairs of repair sites, a total of 79 AsiSI clusters that contain a previously annotated top site was included. The theoretical number of pairs is defined as 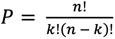, where *n* is the number of AsiSI clusters (79) and *k = 2*, was tested for the presence of DSB-induced (+4OHT) 3D contacts in the logFC Hi-C matrix described above. The mean logFC was calculated within a window of 50-kb surrounding the pixel where both cluster geometric means of the pair meet in the Hi-C matrix. Based on the distribution of mean logFC values across the 3081 pairs, a threshold was set at ≥0.2 to discriminate contacting from not-contacting pairs (Extended Data Figure 4a). Pairs of sites less than 2 Mb apart in linear genomic distance were excluded from the analysis.

#### Analysis of repair coordination

Analysis of repair coordination across single cells is largely categorized in two: 1) based on the binary MSR segment calls, that were used for both pairwise and multi-way similarity metrics (Fig. 4e-g, Extended Data Fig. 4d-e) and 2) based on quantitative depth-normalized repair signal, the Pearson’s coefficient was calculated to evaluate pairwise coordination. The details of both approaches are described below.

##### Coordination of binary MSR repair segments

To measure coordination of binary repair signal, MSR segments overlapping the AsiSI cluster geometric mean were used to determine whether a cluster is repaired in a given cell. A total of 79 AsiSI clusters that contain a previously annotated top site was included. The resulting *m × n* matrix is defined as *M = (a_ij_)*, where *i* is a cell and *j* is an AsiSI cluster. Pairwise distances were calculated between all pairs of clusters using the Jaccard similarity index, defined as the size of the intersection over the size of the union: 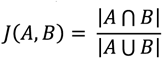 where *A* and *B* represent any pair of clusters in *M*.

For multi-way coordination analysis of repair hubs (*i.e.*, ≥3 contacting AsiSI clusters), we applied a multiple-site extension of the Sørensen-Dice similarity index on binary matrix *M* ^51^. For a detailed justification and explanation of the Sørensen-Dice multi-way similarity metric, see ^51, 57, 58^. We briefly describe the way the metric was used here, below. The multiple-site similarity index for any number of *T* AsiSI clusters can be formulated as

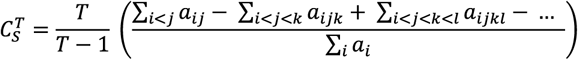

where *a_i_* is the number of cells in which cluster *A_i_* is repaired, *a_ij_* the number of cells that share repair of cluster *A_i_* and *A_j_*, and *a_ijk_* the number of cells that share repair of cluster *A_i_*, *A_j_* and *A_k_*, and so on. In case of *T = 2* the outcome would reflect the definition of the original pairwise Sørensen-Dice similarity index. We wrote a Python implementation of the `betapart` R package ^60^ to compute the multiple-site Sørensen similarity index on a subset of matrix *M* defined as: 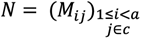 where *c* is a vector of AsiSI cluster indices of any size in *M*. Our `pyBetapart` function is available on GitHub / Zenodo.

##### Coordination of quantitative repair signal

Coordinated behavior of quantitative repair signal for the analysis presented in Figure 3f and (Extended Data) Figure 4 was computed as follows. For intra-chromosomal coordination maps (Figure 3f), the single-cell data was first binned at 20-kb resolution, RPKM-normalized and smoothed using a Gaussian kernel (SD = 4 × 10^4^). For inter-chromosomal coordination maps ((Extended Data) Figure 4), data was binned at 40 kb, RPKM-normalized and smoothed (SD = 4 × 10^5^). For all possible pairs of bins surrounding the AsiSI cluster geometric mean *i* and *j*, repair coordination was calculated as the Pearson’s correlation coefficient between the vectors *b* and *b* that contain the RPKM values of bin *i* and *j* across the single cells.

##### Identifying contacting hubs

The identification of hubs of contacting AsiSI clusters builds on the pair identification described above, but is elaborated by walking through Hi-C matrix *M*^2^ (Extended Data Figure 4g), testing all theoretical combinations of hubs with defined sizes for Hi-C contacts. The hub identification function for any number of *k* AsiSI clusters included in the hub can be written as 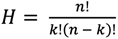 where *n* is the total number of AsiSI clusters included in *M*^2^. All theoretical combinations of hubs were called as contacting hub if all AsiSI clusters within *H* contact each other (Hi-C logFC ≥ 0.2). Our hub identification function written in Python is available on GitHub/ Zenodo.

##### Triplet categorization

For the analysis of (in)complete triplets (Figure 4), we built on our pairwise identification described above. We first determined all combinations of three among a total of 79 AsiSI clusters that contain a previously annotated top site. Theoretical triplets *T* were tested for the presence of Hi-C based contacts (logFC ≥ 0.2) and called “complete” in case all three sites contact each other *(AB, AC, BC)* or “incomplete” if one pair within the triplet does not contact while both other pairs do *(AB, , BC)*.

##### Conditioning of cells for triplet analysis

For triplet analysis presented in Figure 4f-g, cells were stratified by the presence of a binary MSR segment on *both* AsiSI cluster *A and B* (*i.e.*, “AB co-repair” or “synchronous AB”). Cells with an MSR segment on *either A or B* were used for “no AB co-repair” or “asynchronous AB”. In these two conditions of cells, the pairwise coordination of *BC* was measured using the Jaccard similarity index as described above.

##### Coordination Z-score normalization by matrix permutation

Two potential problems are posed on our coordination analysis that might introduce undesirable bias: 1) typical sparsity of single-cell data results in non-uniformly distributed signal dropout and 2) binary similarity metrics can be sensitive to site prominence that differs between sites, but which does not reflect coordination. To solve both problems, we applied a previously described algorithm ^73^, to randomize the abovementioned presence-absence matrix *M n*-times (*n* = 100), without altering row and column totals. The resulting randomized matrices were used to Z-score normalize binary coordination metrics, which can be written as 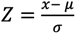, where *x* is the similarity score of the observed, *μ* the mean similarity, and *σ* the standard deviation of the random controls.

#### Multi-way contact triplet analysis in Tri-C data

To measure the presence or absence of multi-way contacts of triplets, we devised an association analysis inspired by previous work ^74^. As described above, a triplet is defined as three AsiSI clusters (A, B and C) that are all observed to contact in the pairwise Hi-C data (AB, AC, BC). In our design, the experimental Tri-C viewpoint (VP) captures AsiSI cluster A, ensuring that each observed read represents a contact that includes A. To discern if A, B and C all contact in a single hub or rather form mutually exclusive contacts, we computationally positively selected reads containing B and quantified the presence of third interaction partner C. In parallel we did the inverse, negatively selecting reads without B and quantify the presence of C. Negative selection was repeated *n*-times (*n* = 100), to create a statistical background profile (mean ±SD) to which the positively selected profile could be compared. Importantly, the positively selected reads are required to contain B, while in negative selection no such constraints are imposed. The positive profile is therefore effectively generated by smaller reads, where one fewer fragment contributes to each profile. To compensate for this technical artifact, we randomly remove one fragment from each read prior to negative selection. The presence of C is statistically quantified by computing the Z-score between the positively and n-times negatively selected profile as 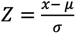, where *x* is the mean of the positively selected Tri-C profile in a defined window surrounding C, *μ* the mean and *σ* the standard deviation in the same window surrounding C, across the *n*-times negatively selected profile. The defined window surrounding C is based on the width of our repair segment enrichment. A positive Z-score indicates that A, B and C preferentially coalesce in a single hub, while a negative Z-score indicates that, although A, B and C all demonstrate pairwise interactions with each other, they preferentially do so in a mutually exclusive manner.

## Notes

https://www.ncbi.nlm.nih.gov/geo/query/acc.cgi?acc=GSE229874

### Competing Interest Statement

The authors have declared no competing interest.

### Summary of Updates

Revised manuscript, including additional multi-factorial and single-molecule experiments. Particularly: Dam&ChIC [Dam-RAD51 & CTCF], SPRITE [validating intercellular variability in genome topology], and Tri-C [validation of higher-order contacts induced by DNA damage].

